# Improved detection of methylation in ancient DNA

**DOI:** 10.1101/2023.10.31.564722

**Authors:** Susanna Sawyer, Pere Gelabert, Benjamin Yakir, Alejandro Llanos Lizcano, Alessandra Sperduti, Luca Bondioli, Olivia Cheronet, Christine Neugebauer-Maresch, Maria Teschler-Nicola, Mario Novak, Ildikó Pap, Ildikó Szikossy, Tamás Hajdu, Eran Meshorer, Liran Carmel, Ron Pinhasi

## Abstract

Reconstructing premortem DNA methylation levels in ancient DNA (aDNA) has led to breakthrough studies such as the prediction of anatomical features of the Denisovan, as well as the castration status of ancient horses. These studies relied on computationally inferring methylation levels from damage signals in naturally deaminated cytosines. Because of statistical constraints, this inference requires high-coverage sequencing, and is thus not only expensive but also restricted to samples with exceptional DNA preservation. Instead, a method to directly measure methylation levels in aDNA, as exists in modern DNA samples, would open the door to a more thorough and cost effective ability to study ancient DNA methylation. We have tested two methods for direct methylation measurement developed for modern DNA based on either bisulfite or enzymatic methylation treatments. We find that both methods preserve sufficient DNA yields to allow for methylation measurement. Bisulfite treatment, combined with a single stranded library preparation, shows the least reduction in DNA yields compared to no methylation treatment, as well as the least biases during methylation conversion. In addition, we show that applying bisulfite treatment to ∼0.4-fold coverage sample provides a methylation signal that is comparable to, or even better, than the computationally inferred one. We thus present a method to directly measure methylation in ancient DNA that is cost effective and can be used on a wide variety of ancient samples.

## Introduction

Since the sequencing of the first human genome, the field of genetics and genomics has made great strides in our understanding of ourselves from medical breakthroughs to a deeper resolution of our history and evolution. As our grasp of the complex relationship between genetics and gene expression has deepened, the study of epigenetics has become integral to understanding genomics. Epigenetics encompasses the study of regulation of genes that are controlled by factors that do not change the genetic code directly. This includes histone modifications, non-coding RNAs and modifications to one of the DNA bases, cytosine (C). The modification of Cs is achieved by the addition of a methyl group, a process referred to as cytosine methylation, or DNA methylation. In vertebrates, the methylation of Cs occurs in the context of a C followed by a guanine (G), also referred to as a CpG context [1], and has been shown to be important in gene regulation [2]. The study of methylation in modern genomes has thus become a powerful tool to understand processes such as aging [3], exposure to chemicals like lead [4] and cancer research [5].

One way to measure methylation directly is to use methods that distinguish between methylated and non-methylated Cs (non-mCs). The most common protocol used today to differentiate between these two C-states, is bisulfite treatment, a protocol that treats single stranded DNA with sodium bisulfate, thereby removing the amine group of non-mCs (a process called deamination) but not deaminating the methylated Cs (mCs). The deamination of non-mCs transforms them into uracils (Us), which are subsequently turned into thymines (Ts) during amplification or sequencing resulting in a C to T misincorporation [6]. After sequencing, reads are aligned to an appropriate reference sequence, which can be used to determine if a C to T change occurred (inferred non-mCs) or if the C stays unchanged (inferred mCs). Bisulfite conversion is an effective tool in modern DNA, however it is a destructive process to DNA [7] and is therefore not optimal for ancient samples. Recently, a new protocol has been developed, the NEBNext^®^ Enzymatic Methyl-seq Kit (EMseq), which has been shown to be effective for only picograms of DNA [8]. The EMseq protocol uses a two-step process of oxidation of mCs to protect them and an enzymatic deamination of unprotected non-mCs, which leads to the same C to T vs C to C differentiation between non-mCs and mCs as produced by the bisulfite conversion gold standard.

Ancient DNA (aDNA) is the study of post mortem DNA. It has been successful for up to 1.2 million year old animals to date [9], as well as allowing for major insights in the human past [10–12]. After death, various processes degrade the DNA of the organism, making it harder to extract and sequence. First, DNA is fragmented into ever shorter fragments [13]. This makes aDNA at best very short and often, typically in older samples or samples from harsh environments, lacking sufficient DNA for meaningful analyses. The minute amounts of DNA also make aDNA susceptible to modern DNA contamination, as well as leaving endogenous DNA at the risk of being overwhelmed by DNA from microbial sources that colonize the bones after death [14].

Bisulfite treatment has previously been applied to aDNA samples. One of the first of these studies applied bisulfite treatment to six ∼26,000-year-old steppe bisons from permafrost. The treated samples were then used as templates in a PCR to amplify four retrotransposon elements in an attempt to determine methylation measurements, however only one of the six had an amplifiable product [15]. A second study also tried to amplify specific regions of interest, in this case LINE-1 elements, in 30 Native American samples after bisulfite treatment. They could show that low DNA concentration leads to higher variability and therefore uncertainty in methylation measurements [16]. A more recent study bisulfite treated tissue samples from two 19th century mummies and subsequently determined methylation levels using the Illumina EPIC BeadChip. They had low DNA concentrations which led to low signal intensities, but it was nevertheless possible to correlate the lung tissue with modern lung tissue and assign the known tissue of origin to the expected tissue-specific methylation levels [17]. None of these studies used next generation sequencing technology and thus relied either on targeting long enough ancient fragments to be able to amplify regions of interest, or bead chips designed for modern DNA. Non-bisulfite-based laboratory methods to evaluate methylation in aDNA have also been promoted, such as using non-U amplifying polymerases [18] and Methyl-Binding-Domain (MBD) enrichment. MBD, which can be done directly on extracts and can therefore circumnavigate issues from bisulfite treatment or capture, has been shown to only be effective for aDNA with long fragments and little natural deamination [19].

One heavily studied aDNA characteristic is the natural deamination of Cs in aDNA fragments. These Us are read as Ts after sequencing, resulting in a C to T misincorporation. Once the reads are sequenced, the rate of C to T misincorporations can be measured according to the location in the original fragment [20]. Studies have shown that, not only does the rate of misincorporation increase toward the end of the fragments [20], the rate also increases with the age of the sample [21].

This C to T misincorporation pattern in aDNA has proven to be a powerful tool in the detection of premortem methylation in aDNA. The deamination of non-mCs turns the Cs into Us, however the methyl group added during the methylation of Cs means that the removal of the amine group transforms Cs directly into Ts. aDNA can be treated with a combination of uracil deglycosylase and Endonuclease VIII (USER treatment) to excise uracils and eliminate the C to T signal left by non-mCs leaving a C to T misincorporation signal caused only by the deamination of mCs [22]. Two main bioinformatic methods have been developed that take advantage of USER treatment to infer methylation levels in ancient DNA, RoAM [23] and DamMet [24]. RoAM has allowed for the study of methylation in Neanderthals and Denisovans, a sister group to Neanderthals. It has led to insights into morphology, brain disorders and possibly even diet [23,25–27]. DamMet was recently used to infer a methylation clock and castration status in ancient horses [28]. As the main source for C to T misincorporation is at the first and last few bases of a fragment, the overall rate of C to T misincorporation due to methylation is small, and accurate reconstruction of the premortem methylation requires a high coverage of the genome of at least 15x on average [29]. This restricts this type of analysis to well preserved aDNA samples subjected to high-coverage shotgun sequencing or targeted capture of specific regions. Additionally, producing high-coverage ancient genomes is expensive in terms of capture and sequencing costs. Furthermore, methylation is measured in windows of consecutive CpG positions, lacking the ability to provide base-pair resolution, as well as only detecting the more common 5-Methylcytosines but not detecting 5-Hydroxymethylcytosines.

A direct examination of methylation through a methylation conversion treatment in aDNA would be invaluable to enable the study of more degraded samples and to measure methylation directly at positions instead of inferring methylation states. Here we examine two different methylation treatments in ancient samples: bisulfite treatment and the EMseq method. We find that the EMseq method, which we hypothesized would be a good candidate for aDNA due to its success with low input amounts, needs further optimizations due to biases that arise in one or both of its two-step conversion process. Meanwhile, bisulfite treatment, in combination with a single stranded library preparation method used for aDNA, provides good performance and could open the door for a cost effective method of direct measurement of aDNA methylation.

## Methods and Results

In order to test various methylation protocols, we chose two samples for which high-coverage data exists as part of the Allen Ancient Genome Diversity Project (https://reich.hms.harvard.edu/ancient-genome-diversity-project). The two samples, Zvej16 (I4438 from [30]) and SP75 (I3957 from [31])), have previously had double stranded libraries produced from cochlea powder DNA extracts with a USER pre-treatment to remove Uracils (Us). The libraries were then shotgun sequenced to a depth of 28.98X and 28.53X, respectively.

Using both the RoAM and DamMet methods, we could produce methylome data for the high-coverage data. Additional extracts were made from the same cochlea bone powder for each of the two samples and were used to apply a range of methylation methods (Tables S1 and S2). Furthermore, we added data from twenty additional samples with varying ranges of DNA preservation for a subset of methods to increase our sample size (Tables S1 and S2). Twelve of these additional samples come from six mummified human remains dating to the 19^th^ century and curated at the Department of Anthropology, Hungarian Natural History Museum (HNHM). The samples include both a cochlea and a molar sample per individual. A lung tissue sample of a different mummified individual from the same crypt was previously bisulfite treated and described in [17]. The other eight samples have produced low-coverage (less than 2-fold) shotgun sequencing data from USER treated libraries as described in [32].

### EMseq in combination with double stranded library methods

As the EMSeq method from NEB (NEBNext^®^ Enzymatic Methyl-seq Kit) has been shown to be effective for methylation conversion of only picograms of DNA [8], we tested the method on our ancient samples. This method entails first converting the DNA fragments into double stranded libraries using the NEB Ultra II library method, with special adapters supplied by the kit that have their non-mCs replaced with mCs. This replacement protects the Cs from the downstream methylation conversions. Next the libraries were methyl converted using the conversion module of the kit. This is a multi-step process, where the mC are oxidized, followed by a denaturation and an enzymatic deamination of the non-mCs. The mCs, being oxidized, are protected from deamination. The final product is an enzymatically deaminated library that has Us instead of non-mCs and has kept the integrity of mCs, which is comparable to the product of bisulfite treatment. This method is subsequently referred to as NEB-EMseq.

As the NEBNext Ultra II library preparation protocol is optimized for modern DNA, we next replaced the library method with a common double stranded library method used in aDNA [33]. We ordered the P5 and P7 adapters with mCs instead of non-mCs and replaced the dNTPs in the final adapter fill-in step with a mixture prepared with d-mCTP instead of d-CTP to assure that the adapters only have mCs. These double stranded libraries, referred to as dslib-EMseq here, were then enzymatically treated with the conversion module part of the NEBNext EM-seq kit to convert non-mCs to Us as described above. See the purple box of Figure 1 for a schematic overview of the double stranded libraries combined with the EMseq methylation conversion.

**Figure 1.**
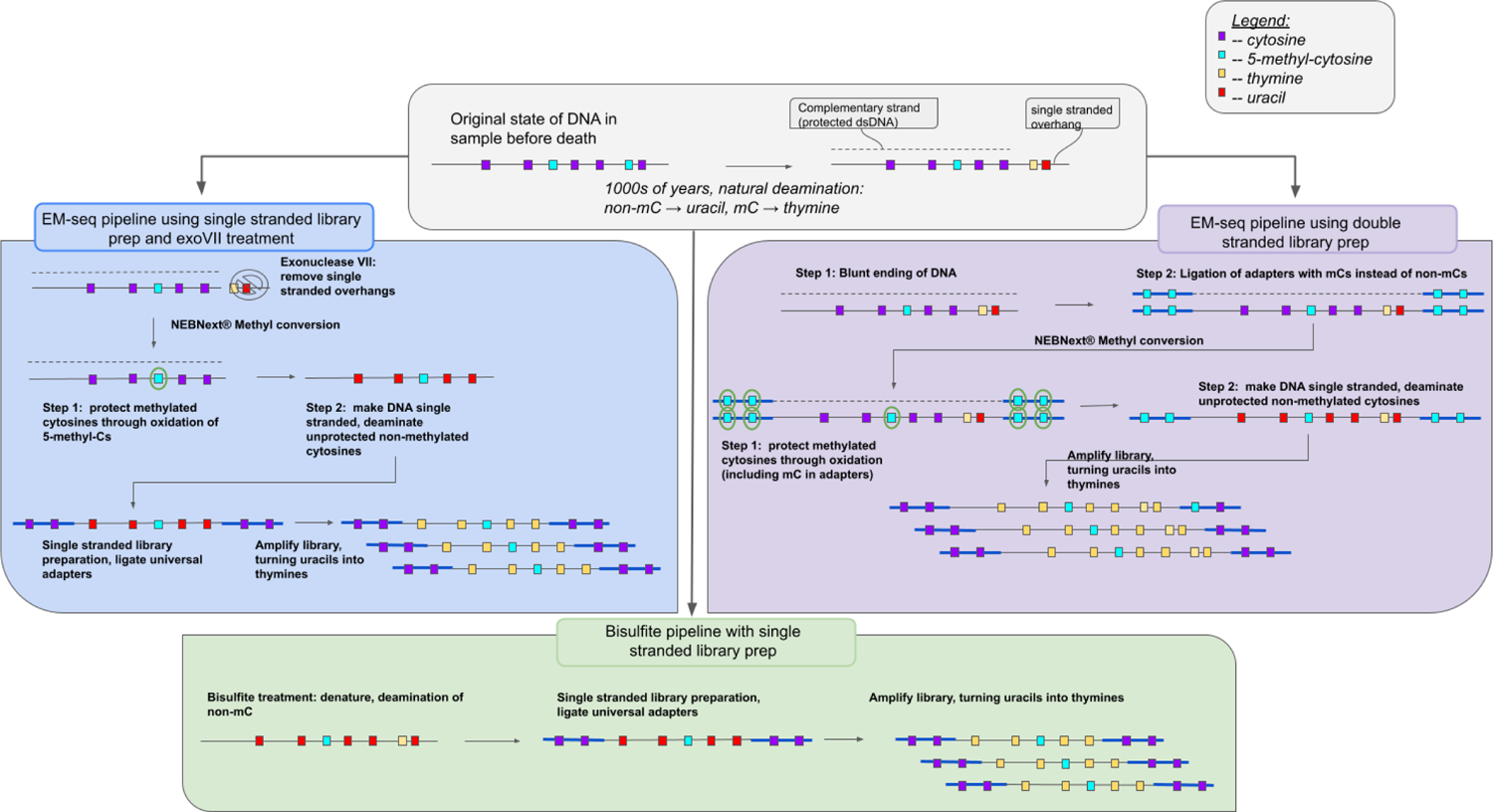
Schematic of the laboratory pipelines tested in this study. DNA extracts were methyl treated either with the EMseq methylation conversion (blue and purple boxes) or with the bisulfite methylation conversion (green box). EMseq was combined either with double stranded library preparation methods (either the NEBNext Ultra II or a double stranded method commonly used in aDNA analyses, purple box), or a single stranded library method developed for aDNA (blue box). The three library prep methods were in addition repeated with a combination of an exonuclease VII pre-treatment, here only shown in the blue box. The bisulfite treatment was only combined with the single stranded library preparation method (green box).

### EMseq in combination with the single stranded library method

As the product of the methyl conversion is single stranded, the double stranded library preparation methods must precede the methyl conversion, necessitating the ordering of adapters with mCs instead of non-mCs. These adapters are expensive due to the number of mCs needed and drive up the cost of the method. Single stranded library preparation methods are more sensitive to aDNA as they convert more fragments into library, and a new single stranded method, the Santa Cruz Reaction (SCR) [34], has been published that is more cost effective and less time consuming than previous aDNA specific single stranded methods [35][36]. We therefore combined the SCR with the EMseq conversion module. However, instead of performing the library preparation first, we first converted the non-mCs into Us enzymatically using the EMseq conversion module, and then followed this conversion with the SCR. This means we can use the SCR protocol and adapters without any changes. We refer to this method as the sslib-EMseq method. See the blue box in Figure 1 for an overview.

### Exonuclease VII (exoVII) treatment

aDNA has a typical C to T misincorporation pattern with an increase in C to T misincorporation the closer the base is to the end of the fragment. This signal is driven by non-mCs and mCs deaminating at the ends of aDNA fragments [20]. As mCs deaminate into Ts directly, they will be indiscernible from the deaminated non-mCs after methylation conversion and could cause a bias in downstream analyses. While deamination has also been shown to happen within fragments due to gaps [37,38], and can happen in blunt end fragment ends, more of the deamination is driven by short single stranded overhangs [38]. We wanted to remove as much of the possible bias of mis-calling mCs due to deamination as possible. To do this, each of the three methods, NEB-EMseq, dslib-EMseq and sslib-EMseq were repeated again with a pre-treatment of the extract with exonucleaseVII (exoVII). The subsequent libraries are referred to as exoVII-NEB-EMseq, exoVII-dslib-EMseq and exoVII-sslib-EMseq respectively.

ExoVII has a 5’ to 3’ as well as a 3’ to 5’ exonuclease activity for single stranded DNA, so treatment of aDNA fragments should fully remove the single stranded overhangs and leave only the double stranded fragments. We tested this with three hybridized oligonucleotides positive controls (see Table S3). One positive control (regular exoVII) had a 60bp strand hybridized to a 40bp strand, allowing for one 10bp overhang on each end. The second control (U-exoVII) had the 60bp strand replaced by a sequence containing Us at the overhangs, while the third (mC-exoVII) had mCs in the overhangs. Each control was then sequenced with and without exoVII treatment. All of the controls showed the exoVII not having full exonuclease activity to blunt end the double stranded DNA, with 3 to 5 bp overhangs remaining despite exoVII treatment (Figure S1). As it has been shown that single stranded overhangs in aDNA are short (1-2 bp) [38], we did not focus on this treatment for further tests as it will require additional optimisation.

### Bisulfite treatment

The last method we applied is bisulfite treatment. We used the EZ DNA Methylation-GoldTM Kit by Zymo Research on each of the extracts, without any changes in the protocol. The single stranded product was then used as a substrate for the SCR [34] to prepare libraries. These libraries are subsequently referred to as bisulfite libraries.

### Indexing, enrichment and sequencing

Each of these libraries were indexed using a double indexing method [39] and a U-tolerant polymerase (Q5U from NEB). After amplification, sslib-EMseq, exoVII-sslib-EMseq and bisulfite libraries of each Zvej16 and SP75 were enriched for the methylome using the methylome capture from Twist Biosciences. We sequenced both the non-captured and captured libraries individually on the Novaseq 6000 system, producing 4-15 million reads per library.

### Positive controls

The NEBNext^®^ Enzymatic Methyl-seq Kit comes with two positive controls to be spiked into each sample before fragmentation: Control DNA CpG methylated pUC19 and Control DNA Unmethylated Lambda. After sequencing, these controls can be aligned to their respective reference genomes and the percent of mC in non-CpG context can be determined (see [8]). As the kit is designed for modern DNA and calls for fragmentation before treatment, the positive controls do not come fragmented and are thus not suitable for aDNA work. Instead, we designed two positive controls that mimic the controls from the kit, but can be examined in more detail after sequencing. Each control is a double stranded 60bp oligonucleotide, one with all 15 Cs methylated (full-methyl-positive) and one with all 15 Cs unmethylated (no-methyl-positive) (Table S3). The controls were spiked into the extract before any treatment or library preparation. For the full-methyl-positive, we expect none of the Cs to be deaminated after treatment. We see that the dslib (with and without exoVII treatment) as well as the non-exoVII treated NEB and the bisulfite treatment have mostly one C deaminated (Figure S2A-B), with only strand 2 showing a slight bias as to position (Figure S2C-D). Meanwhile the other treatments show up to 7 Cs deaminated (Figure S2A). We expect the opposite result for the no-methyl-positive control, with all 15 Cs being deaminated. In strand 1, all treatments cause as few as 7 Cs to be deaminated, with the bisulfite treatment having the highest amount of Cs not deaminated at 15.7% (Figure S3A-B). Again strand 2 shows a more distinct trend of a bias toward which position is less likely to be deaminated, although it is not consistent across treatments (Figure S3C-D).

While these controls cannot be directly compared to the results of the controls from the NEB kit, they do give insights into the effectiveness of methyl conversion in a control situation. Further investigation is needed to understand how CpG context, length or the previous presence of Uracils could affect these controls.

### Effect of methyl treatment on DNA yields

In order to understand how destructive methylation treatment is to aDNA, we first examined the reduction in percent endogenous after methylation conversion in Zvej16 and SP75. When we compare the percent endogenous of each method to DNA extracts made into libraries using the SCR method with no methylation conversion, there is always a reduction of percent endogenous, with the least reduction seen for the bisulfite treatment (Figure 2A, Table S4). As this only represents two samples, we treated additional aDNA samples, from fourteen cochleas and six molars, from various time periods with both the sslib-EMseq and bisulfite methods to increase our sample size to compare percent endogenous. The difference between no methyl treatment and methyl treatment is only significant for the sslib-EMseq method and the exoVII-sslib-EMseq method (Wilcoxson rank sum test, p=0.008 and p=0.002, respectively) but not between no methyl treatment and bisulfite treatment (Wilcoxson rank sum test, p=0.40). For each sample, bisulfite treatment has a higher percent endogenous (0.48 to 26.25%) than the sslib-EMseq method (Table S4) although there is no significant difference between the two treatments (Wilcoxson rank sum test, p=0.056) (Figure 2B). The difference does become significant when comparing the bisulfite treatment to the sslib-EMseq method with an exoVII pretreatment (Wilcoxson rank sum test, p=0.008). A comparison of percent endogenous and ng of DNA used as template for bisulfite treatment shows a slight relationship, however it is not significant (R^2^ = 0.148, p= 0.064 (Figure S5)).

**Figure 2:**
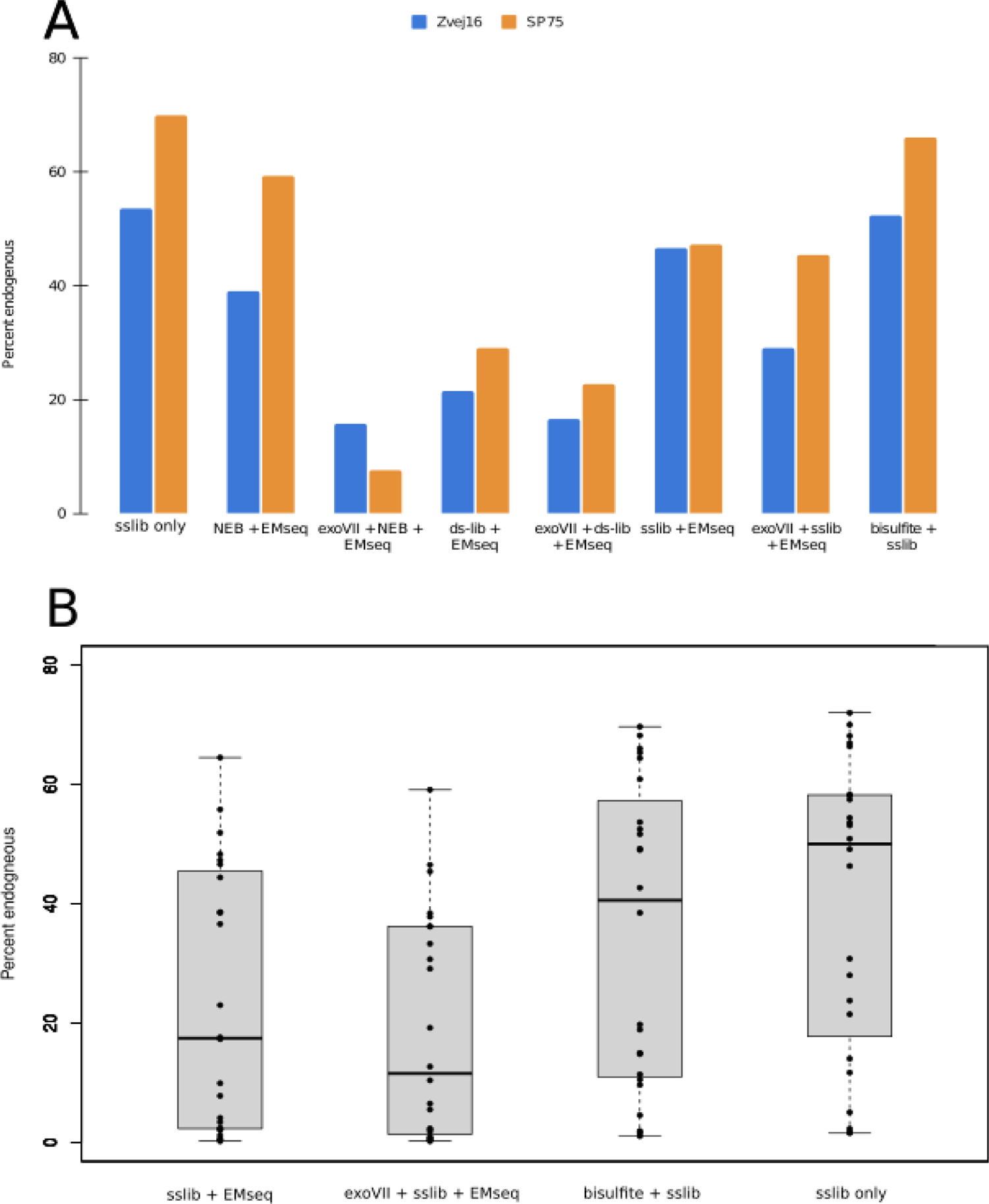
Percent endogenous. Percent endogenous was calculated for each library by dividing the number of mapped reads with a mapping quality of 30 or greater by the raw reads. A) Comparison of the percent endogenous of Zvej16 and SP75 using the single stranded library method with no methyl conversion, to the various methylation conversion and library preparation method combinations. B) Comparison of the percent endogenous of all individuals included in this study using either the EMseq methyl conversion or the bisulfite conversion or neither conversion.

While percent endogenous is an indication of DNA preservation in aDNA, we wanted to understand the effect of the methylation methods on the complexity of the libraries, meaning how many unique library molecules are left after treatment and how much sequencing it takes to start seeing PCR duplicates. Both Zvej16 and SP75 are complex enough samples to produce above 28-fold coverage genomes, so sequencing the libraries deeply enough to see enough PCR duplicates to make inferences on complexity would be prohibitively expensive. Instead, we diluted both extracts to a concentration of 100pg/uL and then used 1uL (corresponding to 100pg) as input for each method, subsequently sequenced each library to produce 10 million reads each. We then subsampled the aligned sequences and checked how many duplicates we see after each subsampling as was done in [10]. For both samples, the non-methyl converted sslib libraries had the highest complexity with the closest fit to infinite complexity (Figure 3). For Zvej16 the bisulfite method has the second highest complexity (Figure 3A) while for SP75, the NEB paired with the EMseq treatment has a higher complexity than bisulfite treatment, however even in this sample, bisulfite has the third highest complexity (Figure 3B). The percent endogenous of the 100pg input is lower than the undiluted extracts (by 2.7% to 51.2%, see Table S4) with again the bisulfite treatment having the highest percent endogenous, even at such low input amounts (Figure S8).

**Figure 3.**
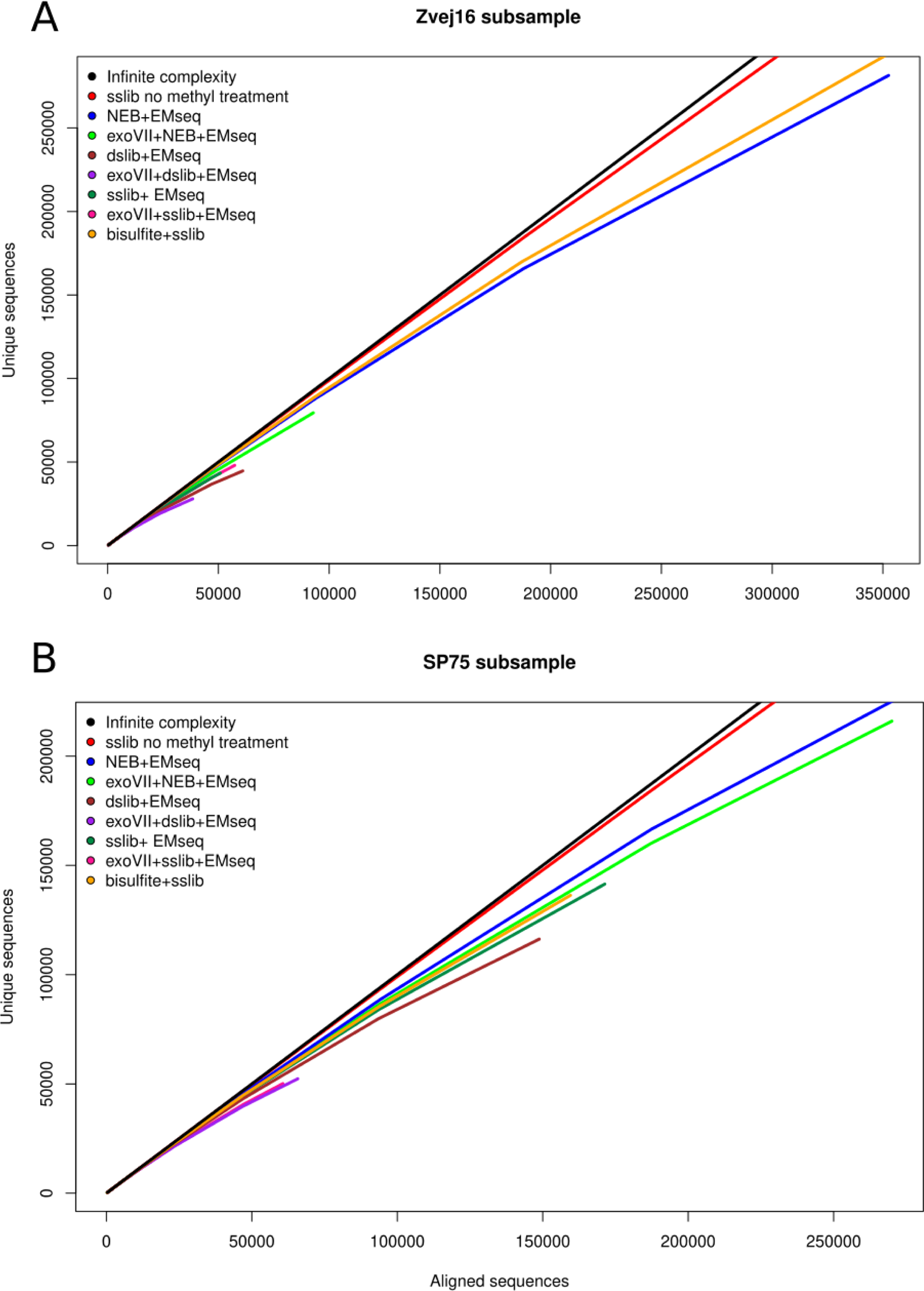
Complexity of reduced extraction input. Subsampling of duplicate removed (unique) sequences versus aligned sequences. The black line represents infinite complexity. Subsampling was done of both Zvej16 (A) and SP75 (B) of 100pg input into various combinations of methyl treatment and library preparation methods.

### Potential biases introduced during methyl treatment

To assess the efficiency of each method in converting non-mCs to Us while not converting mCs to Us, we first examined the percentage of mCs detected bioinformatically in CpG contexts and non-CpG contexts. In mammals, we expect the number of mCs in non-CpG contexts to be low as methylation outside CpG is rare in mammals [40]. Previous studies have found the percentage to be below 0.6% [8]. Meanwhile the average percent of mCs in CpG context is around 70-80% [41]. When examining the CpG contexts of Zvej16 and SP75, all methods involving the EMseq method, except the exoVII-sslib-EMseq method, have above 1% of the Cs in non-CpG contexts defined as mCs (Figure 4, Table S6). This is most likely caused by an issue in the second step of the methylation conversion of the EMseq kit, where non-mCs are not being deaminated and are thus remaining Cs and are subsequently defined as mCs bioinformatically. Interestingly, the NEB method with the exoVII pre-treatment has 52.6-53.4% of mC in CpG context (Table S5), indicating that for this method there is potentially an additional issue with not oxidizing mCs, so they are deaminated in the following step of the conversion module and are thus defined as non-mCs bioinformatically. The bisulfite conversion has the lowest mC percentage in non-CpG context (0.7-0.8%), as well as having a 71.4-74.7% mC in CpG context. As both the exoVII-sslib-EMseq and bisulfite method had the lowest mC in non-CpG context, we wanted to examine if this result is also observed in additional samples. Thus we tested the exoVII-sslib-EMseq, sslib-EMseq and bisulfite libraries from the additional aDNA samples (Table S2) for mC in various contexts and found that the exoVII-sslib-EMseq is not effective at reducing mC in non-CpG contexts for every sample. In fact, two samples, Vác 179 and 11118, see a percentage of mC in non-CpG contexts over 30%. Meanwhile, bisulfite treatment shows a consistent percent of mC in non-CpG context at or below 2.5% for all but three samples tested (Figure 4B). These samples also have some of the lowest percent endogenous as we find that percent endogenous correlates significantly with percentage of mCs in non-CpG context (Figure S6). The difference between the EMseq and bisulfite, as well as EMseq combined with exonuclease VII treatment and bisulfite treatment are significant (Wilcoxson rank sum test, p=4.3e-08 and p=5.9e-07, respectively) (Figure 4B). Interestingly, when examining this result for the 100pg input for Zvej16 and SP75, we see a percentage of below 4% for mCs in non-CpG contexts for all treatments except using the dslib-method (Figure S6). This suggests that low input amounts of DNA are not the only cause for the high mC percent in non-CpG contexts in the additional samples.

**Figure 4.**
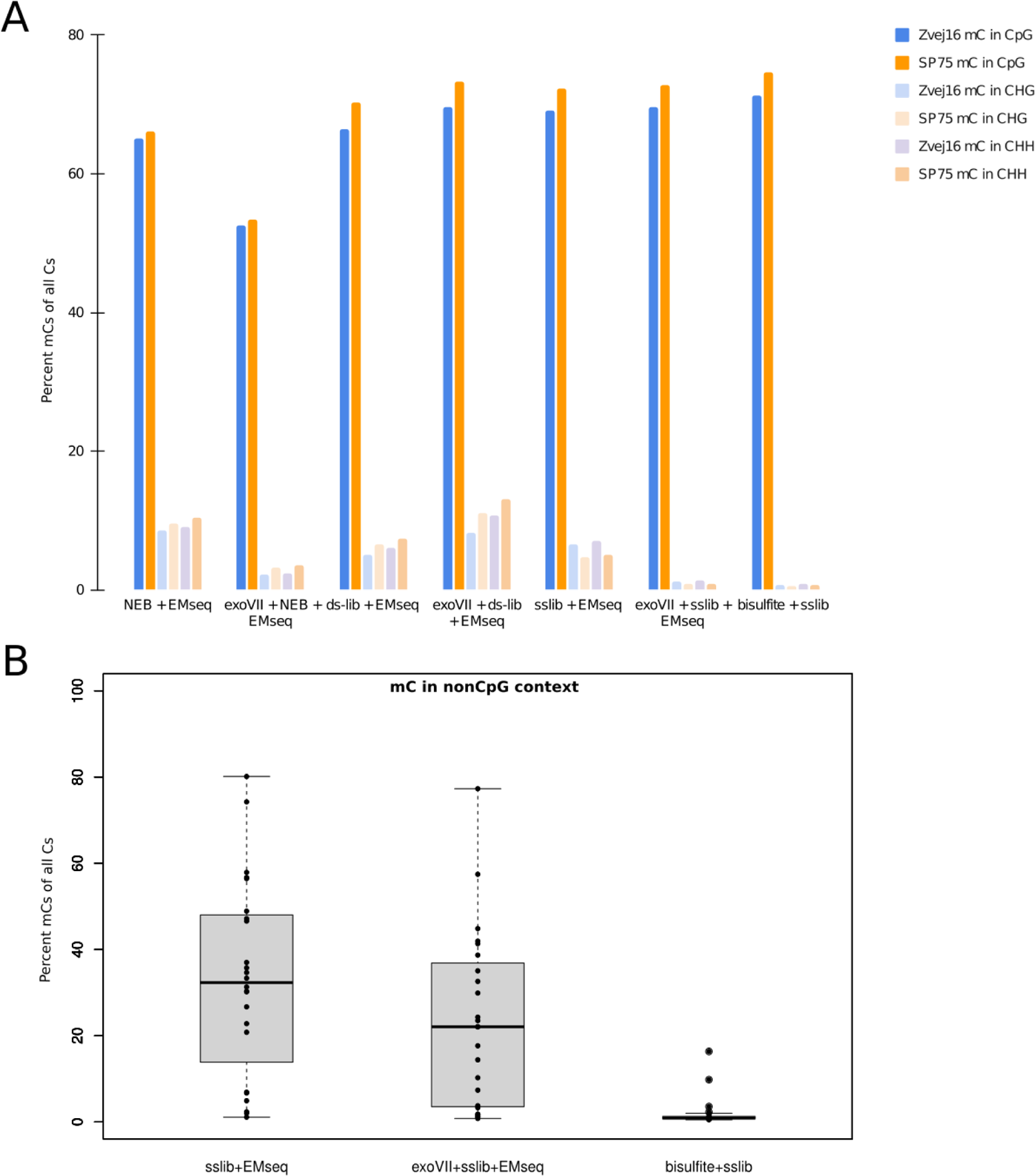
The percentage of methylated Cs of all Cs in various C-contexts. A) Of Zvej16 and SP75 using various methylation treatments in both a CpG context (darker colors) and non-CpG contexts. H refers to a base other than G. B) Of all samples used in this study comparing the EMseq method with and without exonuclease VII treatment in combination with the single stranded library method as well as bisulfite treatment. This comparison is restricted to non-CpG contexts. The two outliers in the bisulfite treatment data are Vác 193 Molar and Vác 164 Petrous.

To examine additional bias sources, we calculated the percentage of mC within and outside of CpG positions of CpG islands. CpG islands are regions of DNA characterized by a high frequency of CpG sites and are often associated with gene regulatory elements. The methylation of Cs in CpG islands is expected to be low, even in CpG context [1]. We see 42.2-83.9 percent higher non-mCs than in mCs in CpG islands, consistent with a reduction in mCs in CpG context in CpG islands. However, the values are variable, with the exoVII-NEB-EMseq having the lowest and bisulfite libraries having the highest percent of mCs in CpG islands (Table S7).

A third bias we examined is the effect of each treatment on read lengths. When comparing the length distributions of various treatments and the single stranded library method without methylation conversion, we find that all methylation conversion methods bias toward longer fragments. The least biases occur in the treatments involving the NEB library preparation method and the EMseq method combined with the single stranded library method and exoVII pre-treatment (Figure S4). This indicates that methylation treatment, be it EMseq or bisulfite, is the most destructive to short aDNA fragments and would be least effective for samples with extremely short fragments.

### Comparison to modern bone and high coverage data

Both the sslib-EMseq and the bisulfite libraries for Zvej16 and SP75 were sequenced further to deeper coverages. We achieved a coverage of 0.46 and 0.61-fold coverage for EMseq for Zvej16 and SP75, respectively, while the bisulfite treatment resulted in a coverage of 0.27 and 0.29-fold coverage for Zvej16 and SP75, respectively. In addition, we captured each of these four libraries using the methylome capture kit from Twist Biosciences and sequenced the capture for another 2.8-14.6 million reads. These sequencing reads were then analyzed with Bismark [42]to calculate a measure of methylation percentage at each position covered. We compared these percentages to both calculated beta values (methylation levels per position) and inferred methylation levels. To simplify comparisons, we subsequently refer to both inferred and calculated methylation levels as beta values. In addition, DNA methylation levels were multiplied by 100 to be comparable to the percentage outputs of Bismark.

As DNA methylation patterns are tissue specific, we compared the beta values of ancient bone samples with modern osteoblast methylation data produced from a 30-fold coverage bisulfite treated sample [43]. In addition, we compared our data to previously described bioinformatic methods to infer methylation [23,24] by inferring beta values from both Zvej16 and SP75 using the high-coverage non-methyl treated and USER treated data from the Allen Ancient Genome Diversity Project. Finally, we examined the inferred beta values of both samples with DamMet using the high coverage data but downsampled to 0.5-fold, 1-fold and 5-fold coverage to better compare to our <1-fold coverage data using the bisulfite and EMseq methods.

After segmenting the data based on the osteoblast beta values, we compared the variation in the histograms of frequency of beta values for each of our comparisons (Figure 5, see Figures S10-S28 for individual histograms). The osteoblast data has 80% of its beta values falling either below 20 or between 71 and 90, in a bimodal distribution of values (Figure 5, Figure S10, Table S8). Both the RoAM beta inference, the low-coverage shotgun data and methylome capture of Zvej16 and SP75 show a similar distribution. An exception is the low-coverage data from SP75 using the EMseq method, where a higher frequency of beta values fall into the 61-70 bin than the 81-90 bin, broadening the peak (Figure 5). The high-coverage DamMet beta inference has a shifted peak with 21-23% of the beta values falling into the 61-70 bin. As the coverage is reduced, the peak also migrates to the left, with the beta inference using DamMet on the 0.5-fold subsampling of the high-coverage data having 20-21% of its beta values fall within the 11-20 bin. In order to understand the relationship between the beta values of our various comparisons, we clustered the beta values using a clustered heatmap (see Supplemental Materials) (Figure 6).

**Figure 5.**
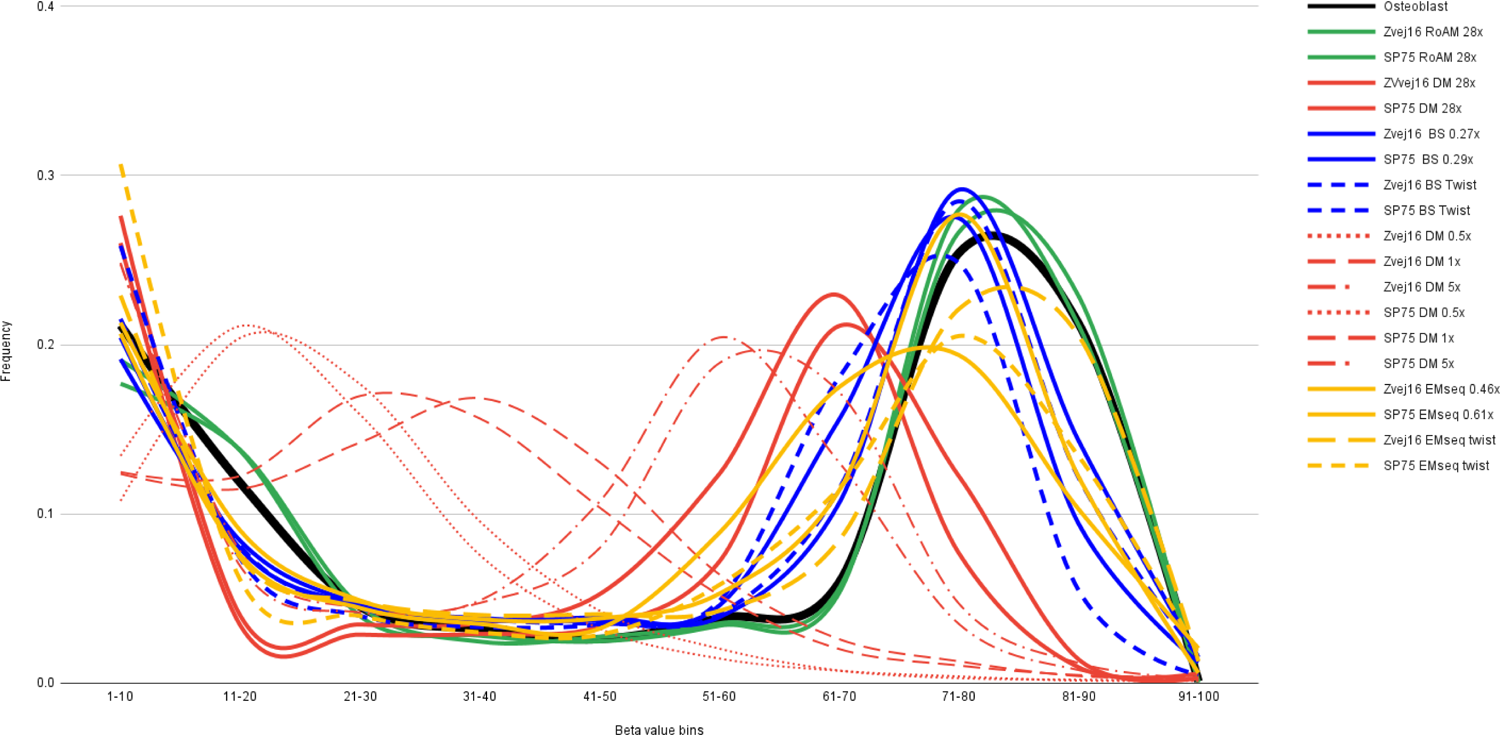
Beta value frequency by bins. The frequency of beta values that fall into bins of 1-100 in 10 increments of chromosome 1. Coverage for shotgun data is shown in the legend. DM = DamMet, BS = bisulfite. Data produced by DamMet and RoAM are using high coverage USER treated data produced of both Zvej16 and SP75 as part of the Allen Ancient Genome Diversity Project. BS and EMseq data of Zvej16 and SP75 were produced from extractions made for this study of the same cochlea as the Allen Ancient Genome Diversity Project and methyl treated either using bisulfite treatment or EMseq treatment in combination with single stranded library preparation. Twist captured data of the methylation treated libraries is also included.

**Figure 6.**
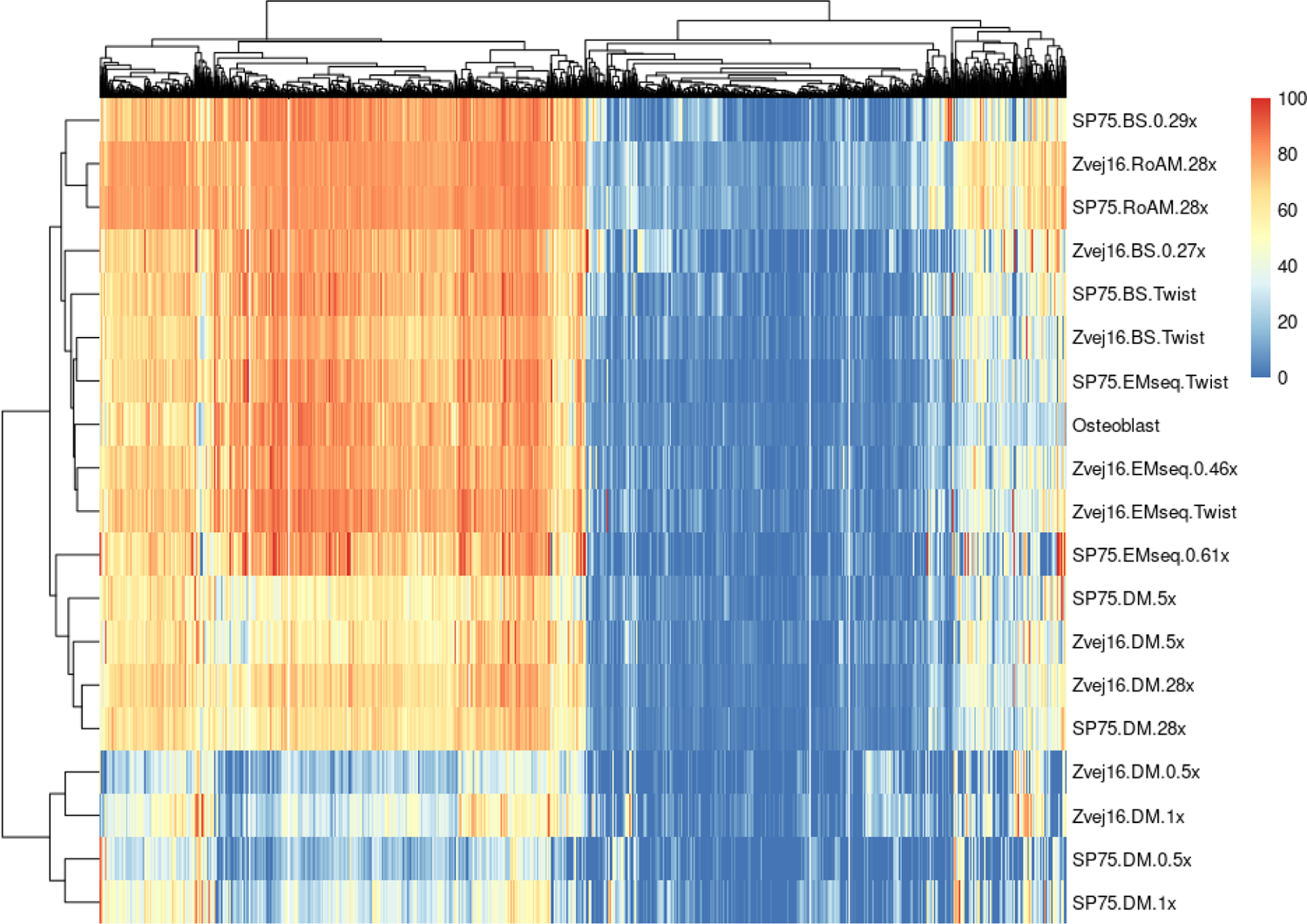
Clustered heatmap of beta values of chromosome 1. The legend shows the beta values with red being a beta value of 100 and blue being a value of 0. DM = DamMet, BS = bisulfite. Data produced by DamMet and RoAM are using high coverage USER treated data produced of both Zvej16 and SP75 as part of the Allen Ancient Genome Diversity Project. BS and EMseq data of Zvej16 and SP75 were produced from extractions made for this study of the same cochlea as the Allen Ancient Genome Diversity Project and methyl treated either using bisulfite treatment or EMseq treatment in combination with single stranded library preparation. Twist captured data of the methylation treated libraries is also included.

The heatmap clusters the DamMet inferred beta values of the 1-fold or lower subsampled high-coverage data into a separate group. The remaining DamMet inferred beta values cluster with the shotgun EM-seq data from SP75, which corresponds to the shifted beta value frequency shift described above. The third main cluster contains the rest of the comparisons, with a subgroup forming containing the RoAM beta value inferences and the SP75 bisulfite treatment shotgun data (Figure 6).

## Discussion

Our results show that ancient DNA can be effectively treated to convert non-mCs to Us to directly measure methylation rates, using two conversion methods developed and perfected for modern DNA. We tried a variation of library preparation methods and conversion methods to understand how effective they are for aDNA. Based on the reduction in percent endogenous, the DNA sequence complexity and the lack of detectable biases in uracil conversion, we find that bisulfite treatment, in combination with an aDNA specific single stranded library preparation method [34], is the most promising pipeline to assess direct methylation rates in aDNA. Even at low-coverage (below 1-fold), this combination of methods performs as well as the previous bioinformatic methods applied to high coverage (over 28-fold) data.

Bisulfite treatment has been used to examine methylation in aDNA in previous studies, usually relying on the use of primers to amplify regions of interest after bisulfite treatment [15,16]. A more recent study [17] examined the methylation signal of a lung tissue sample taken from another mummified individual from the same crypt as the six mummified Vác individuals used in this study (Table S1). This study included an additional mummified sample from Switzerland and used a different approach where they determined methylation based on the Illumina EPIC methylation microarray after bisulfite treatment. They prepared double stranded libraries from non-bisulfite treated extract and had a percent endogenous of 0.65% for the Vác sample after sequencing, indicating a poorly preserved sample. As their data has been produced with a different method and from a different tissue, and methylation has been shown to be heavily tissue dependent [43], and both their data set and ours is small, it is difficult to compare the two datasets.

While bisulfite treatment is the most promising pipeline shown in our study, and has had success in previous aDNA studies, albeit with different methods, additional work should be conducted to optimize the EMseq method for aDNA. The EMseq method is a recent method and has quickly become a popular method to use for methylation treatment in modern DNA. It strives to reduce biases in GC-content and mapping rates seen due to bisulfite treatment [44]. Studies have shown that the EMseq method has fewer biases in GC-content and fragments DNA less than bisulfite treatment [8][45]. However, this is dependent on which bisulfite kit and which library preparation method are used, as bisulfite treatment in combination with a different single stranded library preparation method has been shown to be almost as effective as EMseq, especially for potentially degraded DNA [46].

In our aDNA samples, we show that bisulfite treatment introduces a size bias toward longer fragments, while EMseq combined with the NEB library preparation method reduces the bias toward longer fragments. The biggest hurdle for the NEB-EMseq method is that it introduces biases in non-CpG contexts, most likely occurring due to a reduced efficiency in the second step of the conversion module, where the APOBEC3A enzyme mixture is not deaminating non-mCs effectively. This, in turn, leaves the non-mCs as Cs, which are then assigned as mCs during bioinformatic analyses. This bias is especially surprising, since work on modern plants has shown that the EMseq method reduced this signal compared to bisulfite treatment [45]. One explanation may be that the EMseq method is geared toward a fragment size of 240-290 bp and at least 10ng, thus additional optimization is needed to more effectively deaminate non-mCs in this step. While optimizing the EMseq method for aDNA would be of use to increase the options for methylation conversion in paleoepigenetics, the EMseq method combined with the NEB library preparation method included in the full kit is 3.5 times more expensive than the EZ DNA Methylation-GoldTM Kit we used for bisulfite treatment in combination with the SCR single stranded library preparation method, a cost consideration that is not negligible.

Cs in aDNA deaminate naturally, causing different reactions for mC and non-mCs. Natural deamination of mCs turns them directly into Ts, which in turn causes these mCs to be bioinformatically read as non-mCs. Thus, optimization is needed to address the potential biases that could arise from this signal. In the laboratory, this would, for instance, include additional work on the use of exoVII to completely remove single stranded overhangs. However, deamination has been shown to also occur internally in aDNA fragments in addition to the first few bases of blunt ends [38]. Furthermore, the removal of single stranded overhangs shorten the already short aDNA fragments and, like USER treatment, should be considered carefully on a sample by sample basis as such fragment shortening could significantly affect DNA yields. A better approach would be to explore bioinformatic strategies to estimate and account for this bias. Furthermore, refining current bioinformatic programs designed for modern methylation sequences to be more sensitive to aDNA sequences would also be of value.

We describe here the most comprehensive study of direct methylation detection in aDNA, with a method that can be applied to a multitude of aDNA samples. This method opens the door to a comprehensive study of methylation and subsequent examination of differentially methylated regions in ancient populations, which will provide an additional layer of understanding of various environmental effects such as diet, exposure to diseases and toxins and gestational effects.

## Acknowledgements

We would like to thank Nadin Rohland for helpful feedback on the laboratory process. S.Sa. was funded by the Austrian Science Fund (FWF) M3108-G. P. G. was funded by the Austrian Science Fund (FWF) P–36433. A.L.L was funded by Colfuturo (Fundación para el futuro de Colombia) during his MSc. L.C. and B.Y. were funded by Israel Science Foundation (ISF grant 2436/22). L.C. and E.M. were funded by grant 1001584586 from the knowledge center for forensic DNA Israel by the Israel Ministry of Innovation, Science & Technology. L.C. is the Snyder Granadar chair in Genetics. The biological anthropological work on Vác individuals was supported by the grant of the Hungarian Research, Development and Innovation Office [project numbers: FK128013; TH and IP], the Bolyai Scholarship of the Hungarian Academy of Sciences (TH) and by the ÚNKP-23-5 New National Excellence Program of the Ministry for Culture and Innovation from the source of the National Research, Development and Innovation Fund (TH).

## Author Contributions

S.Sa. conceptualized the study with input from R.P., L.C. and E.M.. S.Sa. and A.L.L. performed laboratory work. S.Sa., P.G. and B.Y. analyzed data. A.S., L.B., O.C., C.N.M., M.T-N., M.N., I.P., I.S. and T.H. excavated and took samples. S.Sa. wrote the paper with input from all authors.

