## Supplemental Materials for "Improved detection of methylation in ancient DNA"

### Supplementary Materials

#### Methods:

##### Laboratory pipelines

***Sample descriptions***

We include a total of 22 samples from 16 individuals in this study (Table S1). Zvej16 (I4438) has been described in [1] and SP75 (I3957) has been described in [2]. In addition, samples 10658, 3543-3547, 2050, 11112 and 11118 have been described in [3] under the names R10658, R3543-R3547, R2050, R11112 and R11118.

During renovation of the White Monks' church of Vác, Hungary in 1994 and 1995, two crypts were uncovered [4,5]. The coffins had been left undisturbed for almost two hundred years and contained the remains of 265 individuals of which six are included in this study (Table S1). Those buried in the crypts were conserved naturally. Mummification was made possible by the unique microclimate and the method of interment. The average temperature of the crypt was low, 8-11° C (46.4 – 51.8 ° F), independent of seasons and external temperature fluctuations. One critical determining factor for the mummification of these individuals was the slight but constant airflow through the two narrow air funnels connecting the undercroft to the outside world. Contributing to the mummification was also the fact that the deceased were placed in coffins made mostly of pine planks and pine wood shavings were placed under the body. The terpenoid content of pine prevented the growth of fungi and bacteria. As a result, the corpses dried up slowly instead of naturally decomposing [6–15].

Based on the descriptions on the coffins and in the parish registers of death and baptisms, the crypt served as a burial place for people living in Vác between 1674 and 1838. The first burial took place in 1731. The archives and inscriptions on the coffins enabled many individuals to be identified by name and occupation – 166 individuals from the 265 are known by name.

The human remains from the crypts are housed in the Department of Anthropology of the Hungarian Natural History Museum, Budapest.

***Bone grinding and DNA extraction***

Cochleas and roots from molars from samples listed in Table S1 were ground in a dedicated clean room as described in [16]. 50 to 100 mg of bone powder was then used to extract DNA using the method from [17] to produce 50uL of DNA extract. Extracts were measured on the Qubit3 with high sensitivity reagents for double stranded DNA to assess the ng/uL concentration. Concentrations of extracts can be found in Table S1. For samples Zvej16 and SP75, an aliquot of each extract was diluted to 100 pg/uL to allow for 100 pg input material for subsampling analyses.

***EMseq***

The NEBNext® Enzymatic Methyl-seq Kit (EMseq) is made up of two relevant parts, one is the library preparation (explained in more detail in the next section) and the other is the methylation conversion module (NEBNext® Enzymatic Methyl-seq Conversion Module -- this module can also be ordered as a stand alone kit). The EMseq conversion was performed as described in the 'Protocol for use with Standard Insert Libraries (370–420 bp)' (NEB #E7120) with the following changes: (1) Neither of the supplied positive controls (Control DNA CpG methylated pUC19 and Control DNA Unmethylated Lambda) was used as they do not come sheared and could not be effectively sheared in our cleanroom. (2) After oxidation, DNA was cleaned up using Minelute columns (Qiagen) by binding to the column with 250uL of buffer PB, washing once with 750uL of wash buffer PE and eluting in 20 uL of buffer EB. (3) Subsequent denaturation was performed by heating the oxidized and cleaned DNA to 95 °C for 3 minutes followed by an ice water bath for 3 minutes. Samples were kept on ice as recommended in the manual. (4) The final deaminated DNA was cleaned using Minelute columns (Qiagen), with 500uL of buffer PB to bind and eluting in either 40uL of EBT (Qiagen buffer EB with 0.05% tween) if combined with either of the double stranded protocols or in 20 uL of EBT if the DNA was then input for the single stranded protocol.

***Bisulfite treatment***

1-10 uL of DNA extract was used as input for bisulfite treatment. The sample was filled to 20 uL using EBT as recommended by the manual. The EZ DNA Methylation-GoldTM Kit by Zymo Research was used for bisulfite treatment. The protocol was followed exactly with no deviations. After elution, the 10 uL of eluate were filled to 20 uL of by adding 10 uL of EBT to prepare for the single stranded library preparation.

***NEBNext UltraII library preparation as part of the EMseq protocol***

1-5 uL of DNA extract was used as input for the NEBNext UltraII library protocol. The protocol and reagents used were part of the NEBNext® Enzymatic Methyl-seq Kit. We used the Protocol for use with Standard Insert Libraries (370–420 bp) (NEB #E7120) with the following changes: (1) We did not include the supplied positive controls (Control DNA CpG methylated pUC19 and Control DNA Unmethylated Lambda). (2) DNA was not fragmented as ancient DNA is already naturally fragmented. (3) The DNA extract was filled up to 50 uL using EBT for the End Prep. (4) Adapter-ligated DNA was cleaned using a Minelute column (Qiagen) by binding the DNA with 468 uL of buffer PB, and eluting in 28 uL of EBT.

***Double stranded library preparation***

1-5uL of DNA extract were used as a template for double stranded library preparation. We followed the protocol outlined in [18], with the following changes: (1) adapters from [18] were ordered to contain mCs instead of Cs to avoid them being deaminated in the methyl conversion. (2) In the adapter fill-in step the regular dNTPs were replaced with dNTPs that contain a d-methyl-CTP instead of a dCTP.

***Single stranded library preparation***

After either EMseq methyl conversion or bisulfite treatment, 20 uL of converted DNA was used as input for the Santa Cruz Reaction as outlined in [19]. No changes were made from the protocol outlined in the supplementary materials of the paper. As the DNA after methyl conversion is single stranded, we could not measure the concentration on the Qubit. Instead we calculated the appropriate SSB, P5 and P7 concentrations using the concentration of the DNA extract before methyl conversion minus 20%.

1-5 uL of DNA extract were also used for each sample to produce single stranded libraries without any methyl conversion.

A description of what sample was treated with which combination of methods can be found in Table S2.

***Exonuclease VII testing and treatment***

In order to reduce potential biases from natural deamination on single stranded overhangs of ancient DNA, we tested using exonuclease VII (NEB M0379S) to remove single stranded overhangs as this enzyme has both 5' - 3' and 3' - 5' exonuclease activity. First we ordered multiple complementary oligonucleotides (Table S3). These were then hybridized in the following combinations: exovii-1 and exovii-2 as a standard test, exovii-3 and exovii-2 to test Us in the overhangs and exovii-4 and exovii-2 to test mCs in the overhangs. Hybridization occurred by combining 10uL each of 20uM concentrated oligonucleotides, 2.5uL of nucleotide-free water and 2.5uL of 10 Oligo hybridization buffer (0.5M NaCl, 0.01M Tris-HCl, 0.001M EDTA) and heating to 95 °C followed by a cooling to 12 °C at -0.1 °C/second. Each hybridized set was then treated with exonuclease VII by adding 10uL of 5X exonuclease VII reaction buffer, 0.2uM of the control oligonucleotide, 0.5uL of 10U/uL exonuclease VII and filling to 50uL with nuclease free water. Both exonuclease VII and non-exonuclease VII treated controls were subsequently made into libraries using the single stranded preparation described above and subsequently indexed and sequenced as described below.

To test exonuclease VII treatment on our samples, Zvej16 and SP75 extracts were treated with exonuclease VII as described above. After MinElute column cleanup, the eluate was then used as input for each of the three library preparation protocols and EMseq combinations.

***Indexing PCR and quality control***

After library preparation and methyl conversion were complete, 1 uL of a 1:40 dilution of each library was measured via qPCR using the IS7 and IS8 primers from [20] and 2X Biozym Blue S´Green (Biozym) master mix. After qPCR the number of copies per sample was measured as well as the number of ideal cycles to amplify without amplifying into plateau (see [20]).

One fourth of each library was then amplified in an indexing PCR using Q5U (NEB) and unique indexes for both the P5 and P7 ends ([20] for list of possible indexes). PCR reactions were performed for various cycles depending on the cycle number calculated during the qPCR. After indexing, the libraries were cleaned using NucleoMag NGS Clean-up and Size Select beads

(Machery-Nagel) at a 1.2-fold concentration. Samples were eluted in 20 uL of EBT, measured on the Qubit3 and the Tapestation.

##### *Twist capture*

After the indexing PCR, Zvej16 and SP75 libraries (see Table S2) were captured using the Twist methylome capture (Twist Biosciences, Twist methylome V1 TE-96341190). First 3 uL of each indexed library was re-amplified using KAPA HiFi 2x MasterMix (Roche) and IS5 and IS6 primers ([18]) for 20 cycles. Reactions were cleaned using the NucleoMag NGS Clean-up and Size Select beads at 1.8-fold concentration. Cleaned amplifications were then dried using a Speedvac for 1 hour at room temperature. After drying, the Twist Biosciences protocol was followed for capture using a modified version for aDNA [21]. After capture, samples were again amplified using KAPA HiFi 2x MasterMix and IS5 and IS6 primers for 20 cycles, cleaned using a 1.8-fold concentration of NucleoMag NGS Clean-up and Size Select beads and measured on the Qubit3 and Tapestation to assess quality.

##### *Positive controls*

As the EMseq protocol comes with positive controls (Control DNA CpG methylated pUC19 and Control DNA Unmethylated Lambda) that are unfragmented, they are not suitable for the cleanroom. Instead we designed our own positive controls: a fully methylated and an unmethylated control (Table S3). Both controls were hybridized as described above in the exoVII section, and then pooled together. Each extract had 0.4 uL of the 0.1uM positive control pool added to the sample input before any treatment took place.

##### *Sequencing and demultiplexing*

Each library was added to sequencing pools to allow for ~5 million sequencing reads, unless deeper sequencing was desired, then up to 15 million reads were sequenced. Pools were sequenced on the NovaSeq 6000 on an XP SP 100 SR cycle flowcell at the Vienna BioCenter Core Facility (VBCF). After sequencing, the VBCF performed basecalling and demultiplexing and sent us demultiplexed fastq files.

#### Bioinformatic pipelines and analyses

##### *Mapping and first quality controls and filters*

Demultiplexed files had adapter sequences and reads shorter than 30 bp removed using cutadapt 4.2 [22]. For mapping we modified the hg19 genome reference using the bismark_genome_preparation package from Bismark [23]. After adapter removal, sequences were mapped to the modified reference genome using Bismark [23] with the single end read parameter. After mapping the percent endogenous of each library was calculated by dividing the number of mapped reads with a mapping quality of 30 or greater by the raw reads (Figure 2 for regular inputs and Figure S8 for subsampled input as well as Table S4). Duplicates were removed using the samtools rmdup package [24]. Duplicate removed reads with a map quality of >=30 were kept for further analyses. Bismark produces multiple output files, one of which calculates the mapping percent, the percentage of mCs in CpG contexts as well as various non-CpG contexts (Figure 4B and S7 for regular inputs and Figure S9 for subsampled input as

well as Tables S5 and S6). Differences in CpG-contexts and percent endogenous were calculated using the Wilcoxon rank sum exact test using R 4.3.1 [25] in Rstudio 2023.06.1. Correlations to produce $R^2$ values were calculated using the correlation function and associated p-values were determined using a linear regression model using R 4.3.1 [25] in Rstudio 2023.06.1 to correlate percent endogenous to ng of input (Figure S5) and percent endogenous to percent of mC in non-CpG contexts (Figure S6).
Non-methyl treated libraries were mapped to the hg19 reference genome, as is standard in aDNA work, using BWA [26] with the parameters -n 0.01 -o 2 -l 16500. After mapping the percent endogenous of each library was calculated by dividing the number of mapped reads with a mapping quality of 30 or greater by the raw reads. Duplicates were removed using the samtools rmdup package [24]. Duplicate removed reads with a map quality of >=30 were kept for further analyses.

***Complexity***
For both Zvej16 and SP75 we input 100pg of extract for each of the treatments. These samples were subsequently shotgun sequenced to produce ~10 million reads. In order to infer complexity, these reduced input samples were filtered as above except duplicates were not removed. Instead the samtools -s option was used to subsample the original bam file reads to produce new bam files with 350 reads and sequentially doubling the read number to 1,500,000 reads. Each new bam file then had duplicates removed and the number of unique (duplicate removed) and total reads were plotted using R 4.3.1 in Rstudio 2023.06.1 using the plot function (Figure 3).

***Exonuclease VII controls***
After sequencing the exonuclease controls, the sequences of both strands of each control (Table S3) were extracted from the raw fastq files and the length distributions of each control was calculated. The frequency of each length was then calculated by dividing the number of reads of each length by the total reads (Figure S1).

***Positive controls***
Positive control reads were extracted from each demultiplexed sample by searching for the correct sequence of both the top and bottom strands (Table S3) in the raw fastq file. The number of reads that match both the top and bottom strands of each of the two positive control types were counted and the percentage of positive controls of the total raw reads was calculated. The values ranged from 0.3-10%. For the full-methyl positive control, each strand has 15 positions of the 60 total that are mCs (and no non-mCs). The number of methylated Cs was counted as well as which position was methylated (Figure S2). For the non-methyl positive, each strand had 15 positions of the 60 total that are non-mCs (and no mCs). Again the number of methylated Cs was counted as well as which position was methylated (Figure S3).

***Length distributions***
The length distributions of the filtered reads (see above) of each treatment possibility (Table S2) of SP75 and Zvej16 were calculated from the final bam files. The frequency of length

distributions was then calculated by dividing the number of reads of each length by the total reads (Figure S4).

***Beta calling***

Osteoblast data: We downloaded the bigwig file from [27] and extracted the beta values as well as chromosome and position data using R 4.3.1 and Rstudio 2023.06.1. These values were merged into a bed file.

RoAM data: We used the RoAM pipeline with default parameters [28] to produce genome-wide reconstruction of premortem DNA methylation of high coverage bam files provided by the Allen Ancient Genome Diversity Project
(Zvej16 (I4438 from [30] at 28-fold coverage and SP75 (I3957 from [31]) at 28-fold coverage).

DamMet data: To calculate F (inference of a beta value), we used DamMet [29] on the high coverage bam files provided by the Allen Ancient Genome Diversity Project
(Zvej16 (I4438 from [30] at 28-fold coverage and SP75 (I3957 from [31]) at 28-fold coverage) using a CpG window size of 20. We then downsampled both samples to 0.5, 1 and 5-fold coverage using the samtools view -s option and re-ran DamMet with the same parameters.

EMseq and bisulfite data: We used the bismark_methylation_extractor package of Bismark [23] to extract the methylation information of each C-position. We use the methylation percentage as a beta value for each position.

As Bismark outputs a percentage, we converted all other beta measurements or approximations to percentages to allow for comparisons.

***CGI controls***

As we expect methylation rates to be lower in CpG islands (CGIs), we calculated the methylation rate in both of these regions and compared them to the total methylation rates for each of our treatments (Table S7). To do this we downloaded the cpgIslandExt track from the UCSC Genome Browser and intersected these positions where either a mC or non-mC (or both) had been called using Bismark using bedtools intersect [32]. We then added the number of mC and non-mCs at each position together and divided the total number of mCs by the total number of mCs plus non-mCs for the original file and the CGI filtered positions.

***Segmentation of data***

We used the R 4.3.1 program methSeg from methylKit [33] to segment the osteoblast file into segments based on their methylation profile with the join.neighbor function turned on to allow neighboring segments that cluster into the same segmentation group to be joined into one segment.

In order to compare beta values for these segments, we used the positions of the segments to filter the other files (RoAM, DamMet, EMseq and bisulfite beta value outputs). Each file was filtered for the positions that fall into the segments determined above and the average beta value per segment was calculated.

***Beta comparisons***

To make comparisons computationally more manageable, we restricted the beta comparisons to chromosome 1. Segmented data from chromosome 1 from each type of treatment and analysis was combined with the segmented osteoblast data into one file. Data was visualized as a histogram using the hist function in R 4.3.1 in Rstudio 2023.06.1 (Figure S10-S28). We further calculated the fraction of beta values that fall into each size bin from 1-100 in bin-increments of 10 (Table S8, Figure 5). Clustered heatmaps of beta values for all samples were made using the pheatmap 1.0.12 function in R 4.3.1 and Rstudio (https://CRAN.R-project.org/package=pheatmap) (Figure 6).

### Supplemental Tables:

Table S1. Samples used in this paper with basic information. Extract concentrations were measured using a Qubit3 high sensitivity double stranded DNA kit. Samples labeled as "Too low" were below the measurable threshold (0.1ng/uL).

| Sample Name | Bone part | Location | Date ([BCE/CE cal 14C*] or [estimated^] ) | Sex | USER treated shotgun seq coverage (no methyl treatment) | ng/uL conc. of extract |
|---|---|---|---|---|---|---|
| SP75 | cochlea | East Altai Mountains, Russia | 1191-1010 BCE* | M | 28.5 | 1.79 |
| Zvej16 | cochlea | Zvejnieki, Latvia | 5462-5220 BCE* | M | 28.9 | 1.83 |
| Vác 179 Petrous (HNHM Inv. No 2009.19.179.) | cochlea | Vác, Hungary | 1800s ^ | F | NA | 0.856 |
| Vác 179 Molar (HNHM Inv. No 2009.19.179.) | First molar | Vác, Hungary | 1800s ^ | F | NA | 0.33 |
| Vác 207 Petrous (HNHM Inv. No 2009.19.207.) | cochlea | Vác, Hungary | 1800s ^ | M | NA | Too low |
| Vác 207 Molar (HNHM Inv. No 2009.19.207.) | First molar | Vác, Hungary | 1800s ^ | M | NA | 0.224 |
| Vác 193 Petrous (HNHM Inv. No 2009.19.193.) | cochlea | Vác, Hungary | 1800s ^ | F | NA | Too low |
| Vác 193 Molar (HNHM Inv. No 2009.19.193.) | Second molar | Vác, Hungary | 1800s ^ | F | NA | 0.186 |

| | | | | | | |
|---|---|---|---|---|---|---|
| Vác 164 Petrous (HNHM Inv. No 2009.19.164.) | cochlea | Vác, Hungary | 1800s ^ | F | NA | Too low |
| Vác 164 Molar (HNHM Inv. No 2009.19.164.) | First molar | Vác, Hungary | 1800s ^ | F | NA | Too low |
| Vác 77 Petrous (HNHM Inv. No 2009.19.77.) | cochlea | Vác, Hungary | 1800s ^ | F | NA | 0.74 |
| Vác 77 Molar (HNHM Inv. No 2009.19.77.) | First molar | Vác, Hungary | 1800s ^ | F | NA | 0.296 |
| Vác 210 Petrous (HNHM Inv. No 2009.19.210.) | cochlea | Vác, Hungary | 1800s ^ | F | NA | 0.736 |
| Vác 210 Molar (HNHM Inv. No 2009.19.210.) | First molar | Vác, Hungary | 1800s ^ | F | NA | 0.304 |
| 10658 | cochlea | Klosterneuburg, Austria | 26-407 CE* | F | 0.8 | 1.06 |
| 3547 | cochlea | Novo Selo, Croatia | 545-597 CE* | F | 0.9 | 0.824 |
| 3543 | cochlea | Gardun, Croatia | 431-600 CE* | M | 0.9 | 0.342 |
| 3544 | cochlea | Gardun, Croatia | 549-600 CE* | M | 1.0 | Too low |
| 3545 | cochlea | Gardun, Croatia | 431-542 CE* | F | 0.8 | 0.33 |
| 2050 | cochlea | Omišalj-Mirine, Croatia | 402-533 CE* | F | 0.9 | 1.05 |
| 11112 | cochlea | Isola Sacra, Italy | 1 -400 CE* | M | 1.1 | Too low |

| | | | | | | |
|---|---|---|---|---|---|---|
| 11118 | cochlea | Isola Sacra, Italy | 1 -400 CE* | F | 1.7 | 1.29 |

Table S2. Samples with various treatments and sequencing strategies

| Sample Name | NEB + EMseq +/- exoVII treatment | dslib + EMseq +/- exoVII treatment | EMseq + sslib +/- exoVII treatment | Bisulfite treatment + sslib | Low-coverage shotgun seq (~5 million reads) | Increased shotgun sequencing (~30-60 million reads) | Twist methylome capture |
|---|---|---|---|---|---|---|---|
| SP75 | yes | yes | yes | yes | yes | yes | yes |
| Zvej16 | yes | yes | yes | yes | yes | yes | yes |
| Vác 179 Petrous | no | no | yes | yes | yes | no | no |
| Vác 179 Molar | no | no | yes | yes | yes | no | no |
| Vác 207 Petrous | no | no | yes | yes | yes | no | no |
| Vác 207 Molar | no | no | yes | yes | yes | no | no |
| Vác 193 Petrous | no | no | yes | yes | yes | no | no |
| Vác 193 Molar | no | no | yes | yes | yes | no | no |
| Vác 164 Petrous | no | no | yes | yes | yes | no | no |
| Vác 164 Molar | no | no | yes | yes | yes | no | no |
| Vác 77 Petrous | no | no | yes | yes | yes | no | no |
| Vác 77 Molar | no | no | yes | yes | yes | no | no |
| Vác 210 Petrous | no | no | yes | yes | yes | no | no |
| Vác 210 Molar | no | no | yes | yes | yes | no | no |

| | | | | | | | |
|---|---|---|---|---|---|---|---|
| 10658 | no | no | yes | yes | yes | no | no |
| 3547 | no | no | yes | yes | yes | no | no |
| 3543 | no | no | yes | yes | yes | no | no |
| 3544 | no | no | yes | yes | yes | no | no |
| 3545 | no | no | yes | yes | yes | no | no |
| 2050 | no | no | yes | yes | yes | no | no |
| 11112 | no | no | yes | yes | yes | no | no |
| 11118 | no | no | yes | yes | yes | no | no |

Table S3. List of oligonucleotides designed for this study

| Oligonucleotide Name | Sequence 5’ - 3’ | Description |
|---|---|---|
| exovii-1 | TCGTCGTTTGGTATGGCTTCATTCAGCTCCG<br>GTTCCCAACGATCAAGGCGAGTTACATGA | Test exonuclease VII cutting ability, strand with overhangs |
| exovii-2 | CGCCTTGATCGTTGGGAACCGGAGCTGAATG<br>AAGCCATAC | Test exonuclease VII cutting ability, strand without overhangs |
| exovii-3 | T**U**GTCGT**U**TGGTATGGCTTCATTCAGCTCCG<br>GTTCCCAACGATCAAGGCGA**U**TTA**U**ATGA | Test exonuclease VII cutting ability including Us, strand with overhangs |
| exovii-4 | T**mC**GTCGTT**mC**GGTATGGCTTCATTCAGCT<br>CCGGTTCCCAACGATCAAGG**mC**GAGTTA**mC**<br>ATGA | Test exonuclease VII cutting ability including mCs, strand with overhangs |
| full-methyl-pos-1 | TmCGTmCGTTTAGTATmCGmCGTmCGTTmC<br>GAmCTTmCGATTmCGmCAAmCGATmCGmC<br>GAmCGAGTTAmCATGA | Positive control containing 15 Cs, all of which are methylated, strand 1 |
| full-methyl-pos-2 | TmCATGTAAmCTmCGTmCGmCGATmCGTT<br>GmCGAATmCGAAGTmCGAAmCGAmCGmC<br>GATAmCTAAAmCGAmCGA | Positive control containing 15 Cs, all of which are methylated, strand 2 |
| no-methyl-pos-1 | TCGTCGTTTGGTATGGCTTCATTCAGCTCCG<br>GTTCCCAACGATCAAGGCGAGTTACATGA | Positive control containing 15 Cs, none of which are methylated, strand 1 |

| | | |
|---|---|---|
| no-methyl-pos-2 | TCATGTAACTCGCCTTGATCGTTGGGAACCGGAGCTGAATGAAGCCATACCAAACGACGA | Positive control containing 15 Cs, none of which are methylated, strand 2 |

Table S4. Percent endogenous. Percent endogenous is calculated by dividing the number of mapped reads with a mapping quality of 30 or greater by the raw reads times 100.

| | sslib (no methyl conversion) | NEB + EMseq | exoVII + NEB + EMseq | ds-lib + EMseq | exoVII + ds-lib + EMseq | ss-lib + EMseq | exoVII + sslib+ EMseq | bisulfite + sslib |
|---|---|---|---|---|---|---|---|---|
| **Zvej16** | 53.6 | 39.1 | 15.7 | 21.5 | 16.6 | 46.6 | 29.1 | 52.5 |
| **SP75** | 70 | 59.4 | 7.6 | 29 | 22.8 | 47.3 | 45.4 | 66 |
| **Zvej16 (100pg input)** | 50.9 | 5.5 | 1.5 | 0.3 | 0.2 | 2.2 | 0.9 | 11.4 |
| **SP75 (100pg input)** | 58.3 | 7.9 | 3.7 | 1.1 | 1 | 4.1 | 2.2 | 14.8 |
| **Vác 179 Petrous** | 54.40 | NA | NA | NA | NA | 38.6 | 38.4 | 51.67 |
| **Vác 179 Molar** | 58.15 | NA | NA | NA | NA | 23.0 | 19.2 | 49.22 |
| **Vác 207 Petrous** | 21.47 | NA | NA | NA | NA | 7.8 | 5.5 | 19.74 |
| **Vác 207 Molar** | 2.29 | NA | NA | NA | NA | 0.2 | 0.2 | 1.91 |
| **Vác 193 Petrous** | 11.70 | NA | NA | NA | NA | 2.4 | 1.8 | 10.51 |
| **Vác 193 Molar** | 5.04 | NA | NA | NA | NA | 1.2 | 0.4 | 4.56 |
| **Vác 164 Petrous** | 1.69 | NA | NA | NA | NA | 0.6 | 0.3 | 1.05 |
| **Vác 164 Molar** | 23.77 | NA | NA | NA | NA | 2.1 | 0.8 | 14.99 |
| **Vác 77 Petrous** | 57.48 | NA | NA | NA | NA | 38.5 | 36.2 | 53.66 |
| **Vác 77 Molar** | 1.56 | NA | NA | NA | NA | 0.5 | 0.6 | 1.57 |
| **Vác 210 Petrous** | 14.02 | NA | NA | NA | NA | 3.4 | 2.3 | 9.63 |
| **Vác 210 Molar** | 72.01 | NA | NA | NA | NA | 51.9 | 46.5 | 65.31 |
| **10658** | 30.78 | NA | NA | NA | NA | 9.9 | 6.5 | 18.90 |
| **3547** | 46.31 | NA | NA | NA | NA | 17.3 | 10.4 | 38.47 |
| **3543** | 53.13 | NA | NA | NA | NA | 48.3 | 37.8 | 64.40 |

| | | | | | | | | |
|---|---|---|---|---|---|---|---|---|
| **3544** | 68.12 | NA | NA | NA | NA | 55.8 | 36.2 | 68.19 |
| **3545** | 66.36 | NA | NA | NA | NA | 44.4 | 33.3 | 60.88 |
| **2050** | 66.92 | NA | NA | NA | NA | 64.5 | 59.1 | 69.67 |
| **11112** | 28.00 | NA | NA | NA | NA | 17.6 | 12.7 | 42.68 |
| **11118** | 49.13 | NA | NA | NA | NA | 36.6 | 30.7 | 49.01 |

Table S5. Percentage of mC in CpG context.

| | **NEB + EMseq** | **exoVII + NEB + EMseq** | **ds-lib + EMseq** | **exoVII + ds-lib + EMseq** | **ss-lib+EMseq** | **exoVII + sslib+ EMseq** | **bisulfite + sslib** |
|---|---|---|---|---|---|---|---|
| **Zvej16** | 65.1 | 52.6 | 66.5 | 69.6 | 69.1 | 69.6 | 71.2 |
| **SP75** | 66.1 | 53.4 | 70.3 | 73.3 | 72.2 | 72.7 | 74.7 |
| **Zvej16 (100pg input)** | 56.3 | 58.7 | 68.7 | 71.8 | 57.6 | 59.9 | 67.6 |
| **SP75 (100pg input)** | 57.7 | 60 | 72.1 | 73.3 | 61 | 64 | 71.6 |
| **Vác 179 Petrous** | NA | NA | NA | NA | 57.2 | 79.8 | 73.9 |
| **Vác 179 Molar** | NA | NA | NA | NA | 82.6 | 85.3 | 75.8 |
| **Vác 207 Petrous** | NA | NA | NA | NA | 55.9 | 78 | 76.9 |
| **Vác 207 Molar** | NA | NA | NA | NA | 55.1 | 50.1 | 46 |
| **Vác 193 Petrous** | NA | NA | NA | NA | 64.8 | 74.7 | 72.4 |
| **Vác 193 Molar** | NA | NA | NA | NA | 64.1 | 59.7 | 68 |
| **Vác 164 Petrous** | NA | NA | NA | NA | 46.9 | 58.4 | 45.7 |
| **Vác 164 Molar** | NA | NA | NA | NA | 55.6 | 48.1 | 58.6 |
| **Vác 77 Petrous** | NA | NA | NA | NA | 55.8 | 78.1 | 75.6 |

| | | | | | | | |
|---|---|---|---|---|---|---|---|
| **Vác 77 Molar** | NA | NA | NA | NA | 79.9 | 82.5 | 58.5 |
| **Vác 210 Petrous** | NA | NA | NA | NA | 71.6 | 75.2 | 74.8 |
| **Vác 210 Molar** | NA | NA | NA | NA | 70 | 82.5 | 76.2 |
| **10658** | NA | NA | NA | NA | 49 | 72.8 | 71.8 |
| **3547** | NA | NA | NA | NA | 47.3 | 68.9 | 69 |
| **3543** | NA | NA | NA | NA | 49.9 | 64.9 | 69.4 |
| **3544** | NA | NA | NA | NA | 41.6 | 74.8 | 72 |
| **3545** | NA | NA | NA | NA | 53.8 | 75.6 | 74.3 |
| **2050** | NA | NA | NA | NA | 52.2 | 74.9 | 72.7 |
| **11112** | NA | NA | NA | NA | 30.2 | 71 | 66.4 |
| **11118** | NA | NA | NA | NA | 71.3 | 79.2 | 75.5 |

Table S6. Percentage of mC in non-CpG context.

| | **NEB + EMseq** | **exoVII + NEB + EMseq** | **ds-lib + EMseq** | **exoVII + ds-lib + EMseq** | **ss-lib+EMseq** | **exoVII + sslib+ EMseq** | **bisulfite + sslib** |
|---|---|---|---|---|---|---|---|
| **Zvej16** | 8.9 | 2.3 | 5.6 | 9.5 | 6.9 | 1.3 | 0.8 |
| **SP75** | 10 | 3.4 | 7 | 12 | 4.9 | 0.8 | 0.7 |
| **Zvej16 (100pg input)** | 1.3 | 1 | 20.1 | 17.6 | 2 | 3.3 | 1.1 |
| **SP75 (100pg input)** | 1.1 | 1 | 16.8 | 14.3 | 1.1 | 1.8 | 1 |
| **Vác 179 Petrous** | NA | NA | NA | NA | 35.75 | 32.6 | 0.7 |
| **Vác 179 Molar** | NA | NA | NA | NA | 74.25 | 57.45 | 0.6 |

| | | | | | | | |
|---|---|---|---|---|---|---|---|
| **Vác 207 Petrous** | NA | NA | NA | NA | 34.65 | 29.9 | 1.15 |
| **Vác 207 Molar** | NA | NA | NA | NA | 56.45 | 44.85 | 3.55 |
| **Vác 193 Petrous** | NA | NA | NA | NA | 47.15 | 22.05 | 1.4 |
| **Vác 193 Molar** | NA | NA | NA | NA | 46.6 | 38.7 | 16.35 |
| **Vác 164 Petrous** | NA | NA | NA | NA | 20.8 | 23.5 | 9.8 |
| **Vác 164 Molar** | NA | NA | NA | NA | 37 | 22.1 | 2.3 |
| **Vác 77 Petrous** | NA | NA | NA | NA | 33.35 | 24.3 | 0.6 |
| **Vác 77 Molar** | NA | NA | NA | NA | 80.15 | 77.3 | 2 |
| **Vác 210 Petrous** | NA | NA | NA | NA | 56.7 | 41.95 | 1 |
| **Vác 210 Molar** | NA | NA | NA | NA | 48.9 | 41.35 | 0.5 |
| **10658** | NA | NA | NA | NA | 31.3 | 14.4 | 0.8 |
| **3547** | NA | NA | NA | NA | 26.7 | 7.35 | 0.65 |
| **3543** | NA | NA | NA | NA | 22.8 | 3.75 | 0.65 |
| **3544** | NA | NA | NA | NA | 6.7 | 1.7 | 0.85 |
| **3545** | NA | NA | NA | NA | 30.3 | 10.25 | 1 |
| **2050** | NA | NA | NA | NA | 30.2 | 17.65 | 0.8 |
| **11112** | NA | NA | NA | NA | 2.35 | 1.2 | 0.75 |
| **11118** | NA | NA | NA | NA | 57.9 | 35.05 | 1.15 |

Table S7. CpG island control. The percentage of methylated and non-methylated Cs is calculated within CpG islands for each treatment.

| Treatment | Zvej16 | | SP75 | |
|---|---|---|---|---|
| | mCs within CGI | non-mCs within CGI | mCs within CGI | non-mCs within CGI |
| NEB+EMseq | 21.0 | 79.0 | 21.4 | 78.6 |
| exoVII+NEB+EMseq | 10.3 | 89.7 | 8.0 | 91.9 |
| dslib+EMseq | 16.4 | 83.6 | 19.3 | 80.7 |
| exoVII+dslib+EMseq | 17.9 | 82.1 | 21.3 | 78.7 |
| sslib+EMseq | 17.9 | 82.1 | 17.4 | 82.6 |
| exoVII+sslib+EMseq | 13.9 | 86.1 | 13.3 | 86.7 |
| bisulfite+sslib | 26.1 | 73.9 | 28.9 | 71.1 |

Table S8. Distribution of beta values per bin. Beta values are binned from 1-100 in bins of 10. Note that beta values of zero are not included. Coverage for shotgun data is shown in the legend. DM = DamMet, BS = bisulfite. Data produced by DamMet and RoAM are using high coverage USER treated data produced of both Zvej16 and SP75 as part of the Allen Ancient Genome Diversity Project. BS and EMseq data of Zvej16 and SP75 were produced from extractions made for this study of the same cochlea as the Allen Ancient Genome Diversity Project and methyl treated either using bisulfite treatment or EMseq treatment in combination with single stranded library preparation. Twist captured data of the methylation treated libraries is also included.

| | **Bins of beta values** | | | | | | | | | |
|---|---|---|---|---|---|---|---|---|---|---|
| Sample | **1-10** | **11-20** | **21-30** | **31-40** | **41-50** | **51-60** | **61-70** | **71-80** | **81-90** | **91-100** |
| Osteoblast | 0.21 | 0.12 | 0.05 | 0.03 | 0.03 | 0.04 | 0.06 | 0.25 | 0.21 | 0.00 |
| Zvej16 RoAM 28x | 0.19 | 0.14 | 0.04 | 0.03 | 0.02 | 0.03 | 0.05 | 0.28 | 0.21 | 0.00 |
| SP75 RoAM 28x | 0.18 | 0.14 | 0.05 | 0.02 | 0.03 | 0.04 | 0.06 | 0.27 | 0.23 | 0.00 |
| ZVvej16 DM 28x | 0.26 | 0.03 | 0.03 | 0.03 | 0.03 | 0.07 | 0.21 | 0.12 | 0.01 | 0.00 |
| SP75 DM 28x | 0.28 | 0.04 | 0.03 | 0.04 | 0.05 | 0.12 | 0.23 | 0.08 | 0.01 | 0.01 |
| Zvej16 BS 0.27x | 0.20 | 0.08 | 0.05 | 0.04 | 0.04 | 0.05 | 0.16 | 0.27 | 0.09 | 0.01 |
| SP75 BS 0.29x | 0.19 | 0.08 | 0.05 | 0.04 | 0.04 | 0.04 | 0.11 | 0.29 | 0.14 | 0.02 |
| Zvej16 BS Twist | 0.26 | 0.07 | 0.04 | 0.03 | 0.03 | 0.05 | 0.18 | 0.25 | 0.05 | 0.00 |
| SP75 BS Twist | 0.22 | 0.08 | 0.04 | 0.04 | 0.04 | 0.04 | 0.11 | 0.28 | 0.12 | 0.02 |
| Zvej16 DM 0.5x | 0.11 | 0.20 | 0.18 | 0.10 | 0.04 | 0.02 | 0.01 | 0.00 | 0.00 | 0.00 |
| Zvej16 DM 1x | 0.12 | 0.11 | 0.14 | 0.17 | 0.13 | 0.06 | 0.03 | 0.01 | 0.00 | 0.00 |
| Zvej16 DM 5x | 0.25 | 0.07 | 0.04 | 0.05 | 0.08 | 0.19 | 0.17 | 0.05 | 0.01 | 0.00 |
| SP75 DM 0.5x | 0.13 | 0.21 | 0.16 | 0.08 | 0.03 | 0.01 | 0.01 | 0.00 | 0.00 | 0.00 |
| SP75 DM 1x | 0.12 | 0.12 | 0.17 | 0.16 | 0.10 | 0.05 | 0.02 | 0.01 | 0.00 | 0.00 |
| SP75 DM 5x | 0.25 | 0.08 | 0.04 | 0.05 | 0.11 | 0.20 | 0.13 | 0.03 | 0.01 | 0.00 |
| Zvej16 EMseq 0.46x | 0.21 | 0.09 | 0.05 | 0.04 | 0.04 | 0.05 | 0.11 | 0.28 | 0.12 | 0.01 |
| SP75 EMseq 0.61x | 0.21 | 0.08 | 0.04 | 0.03 | 0.03 | 0.09 | 0.17 | 0.19 | 0.10 | 0.02 |

| | | | | | | | | | | |
|---|---|---|---|---|---|---|---|---|---|---|
| Zvej16 EMseq twist | 0.23 | 0.07 | 0.05 | 0.04 | 0.04 | 0.04 | 0.08 | 0.22 | 0.21 | 0.01 |
| SP75 EMseq twist | 0.31 | 0.06 | 0.04 | 0.03 | 0.03 | 0.06 | 0.12 | 0.21 | 0.13 | 0.02 |

### Supplemental Figures:

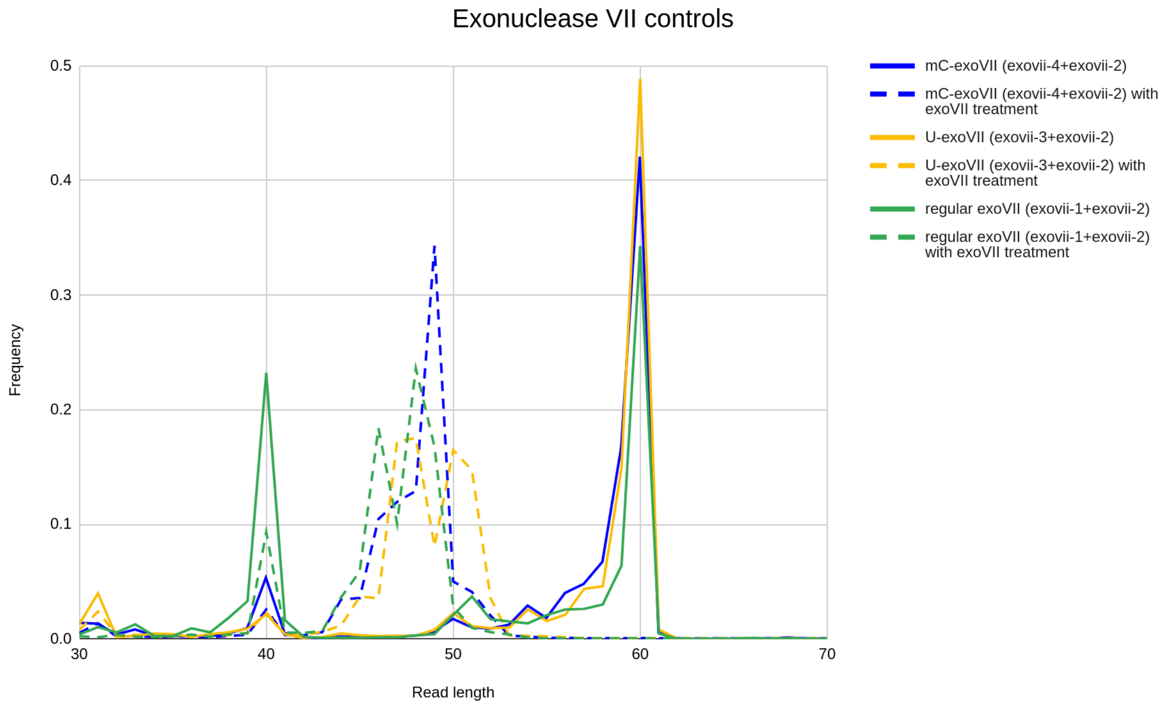


Figure S1. Exonuclease VII controls. In order to understand how well the exonuclease activity of exonuclease VII acts on overhangs, and what may affect the activity, we show the frequency of reads with various read lengths for each type of experiment.

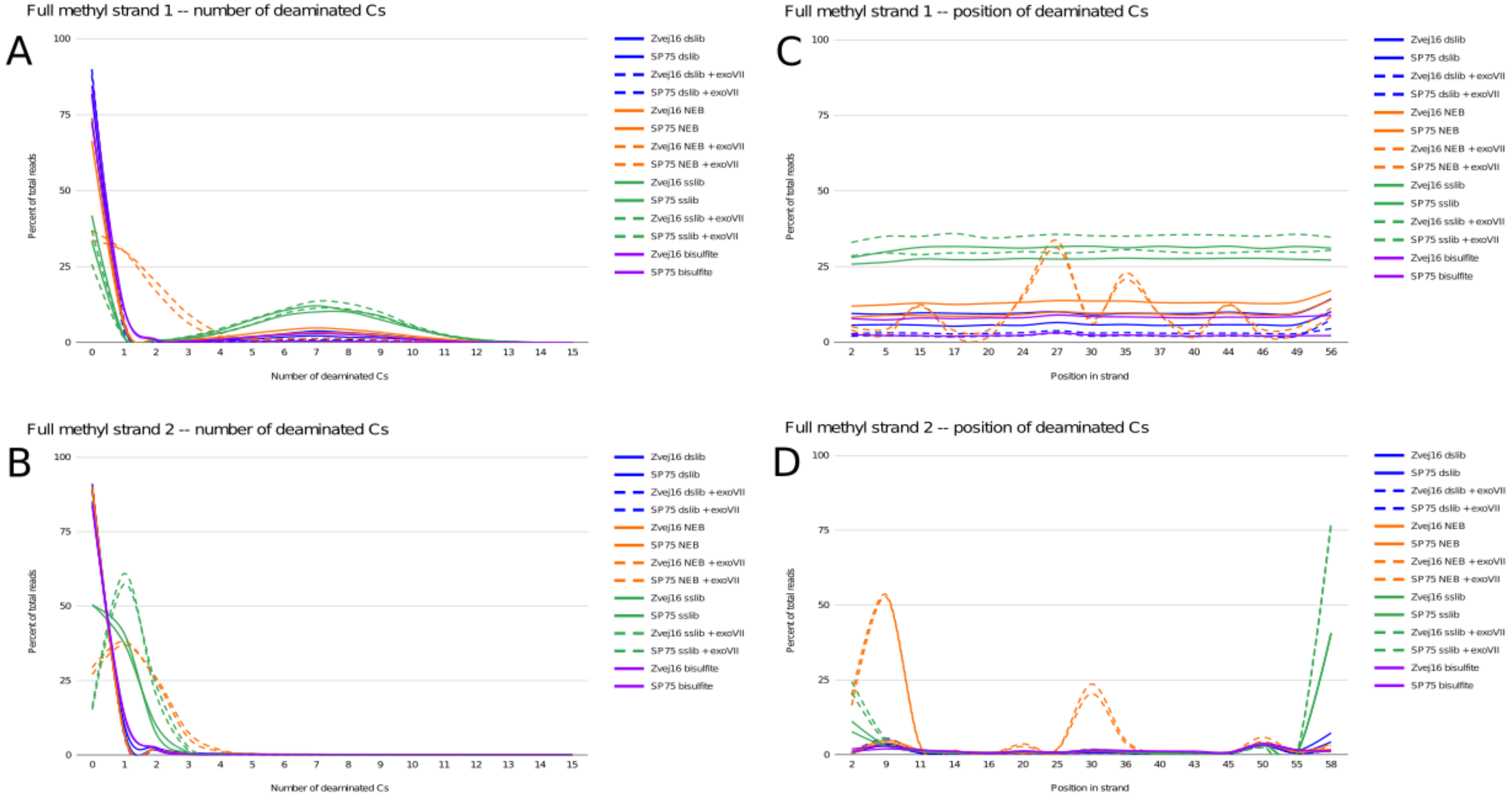


Figure S2. Full methyl positive controls. A) and B) depict the number of Cs that are deaminated in each of the two strands of the positive control. As this control has all 15 Cs in the 60bp control methylated, we expect none of the Cs to be deaminated if the positive control works with perfect efficiency. C) and D) depict the position in the 60bp strand that has a C and is deaminated.

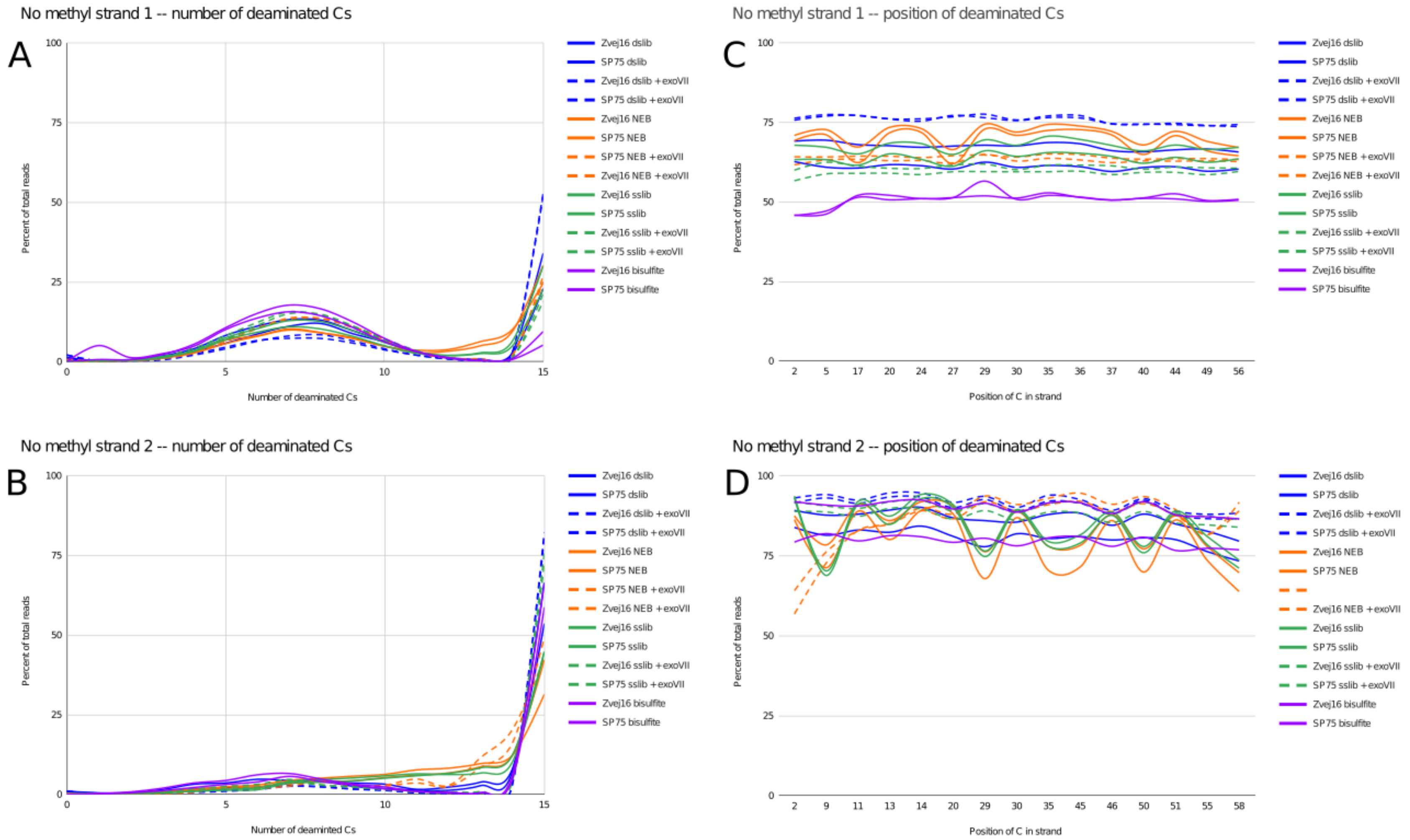


Figure S3. No methyl positive controls. A) and B) depict the number Cs that are deaminated in each of the two strands of the positive control. As this control has none of the 15 Cs in the 60bp control methylated, we expect all of the Cs to be deaminated if the positive control works with perfect efficiency. C) and D) depict the position in the 60bp strand that has a C and is deaminated.

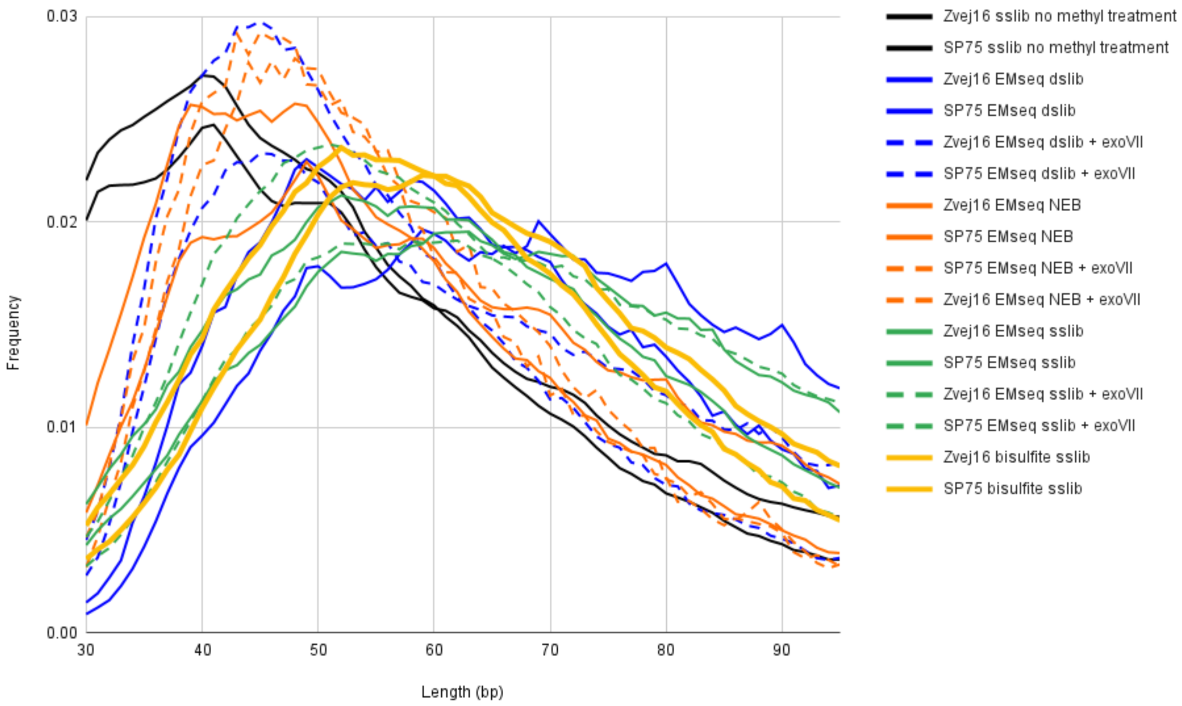


Figure S4. Length distributions of Zvej16 and SP75 after each of the different treatments.

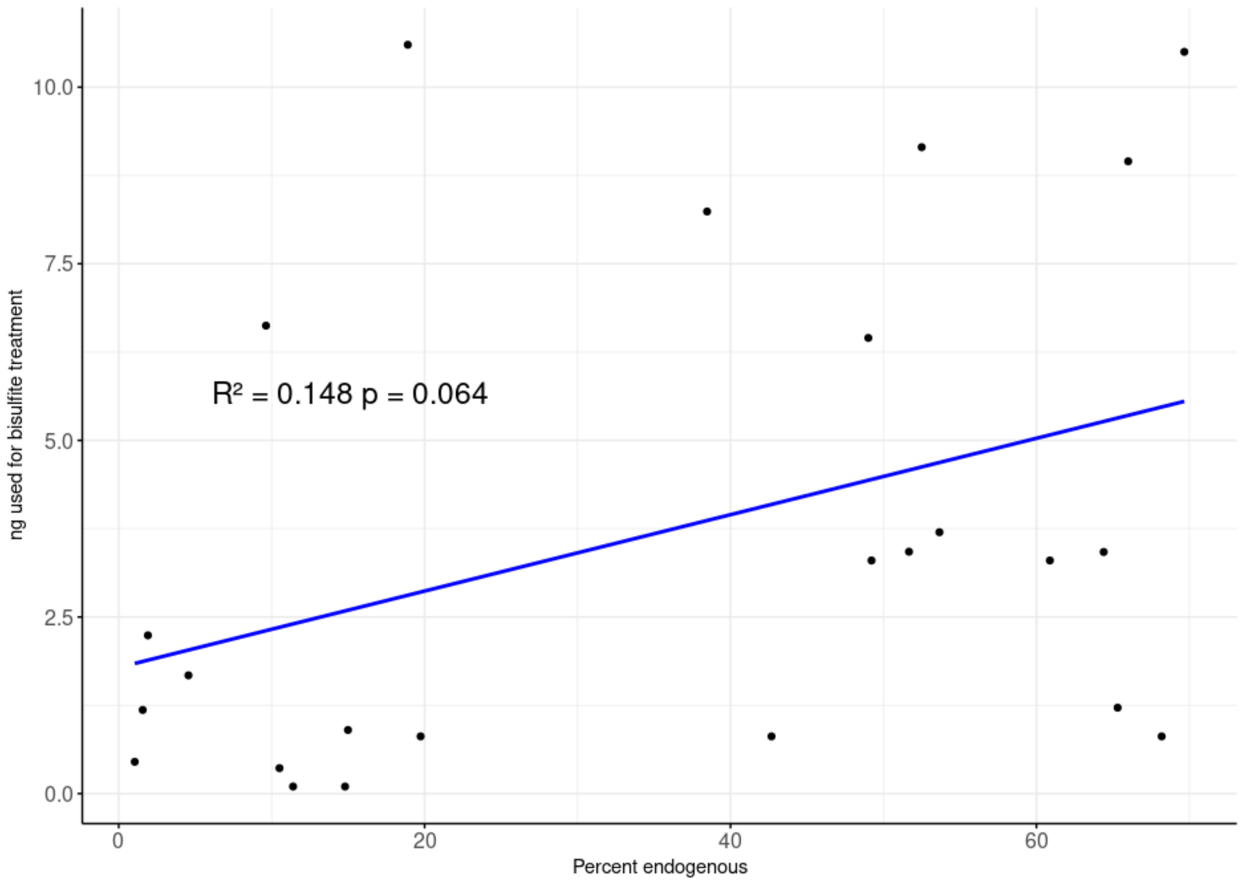


Figure S5. A scatterplot of percent endogenous versus the amount of ng used as input for bisulfite treatment. Each dot represents one of the samples from Table S4.

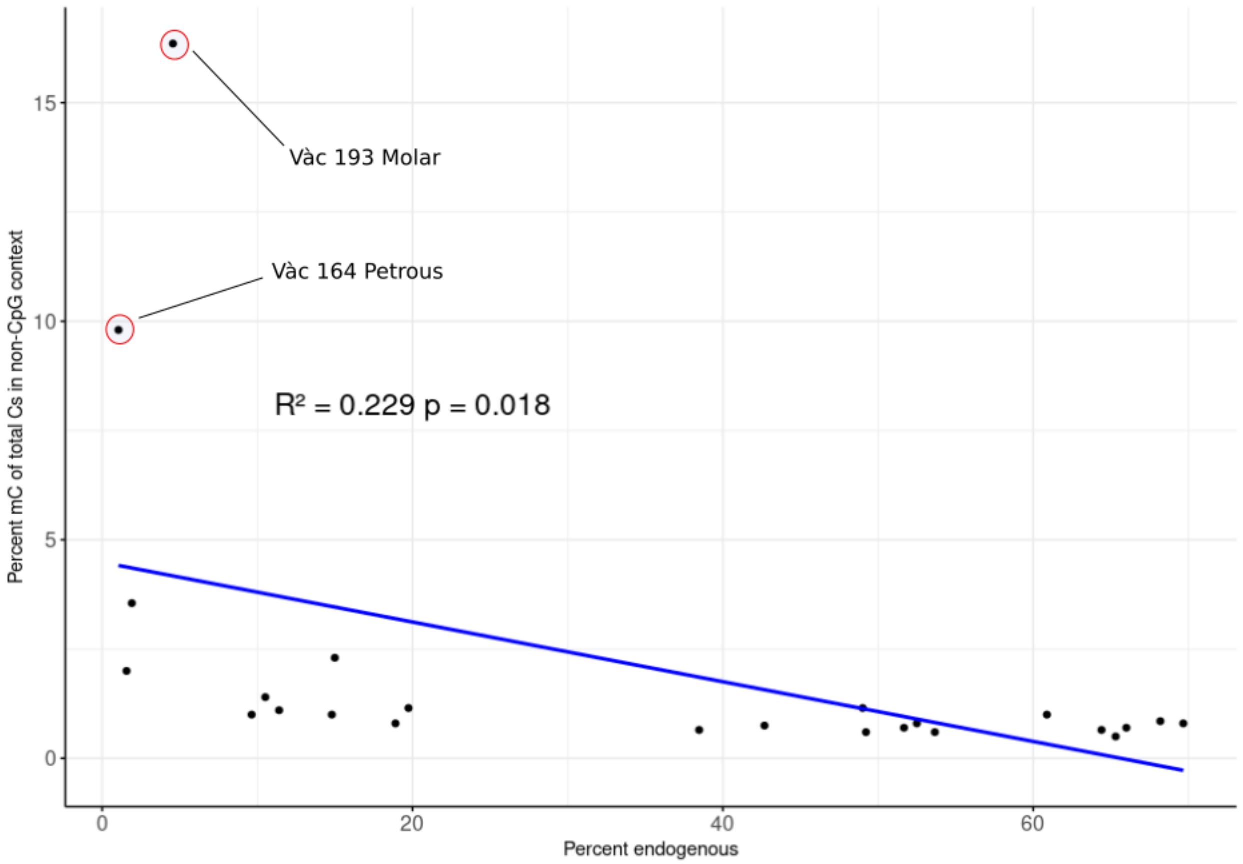


Figure S6. A scatterplot of percent endogenous versus the percentage of mCs of total Cs in a non-CpG context. Each dot represents one of the samples from Table S4. The two samples with the highest percentage of mCs in non-CpG context are shown.

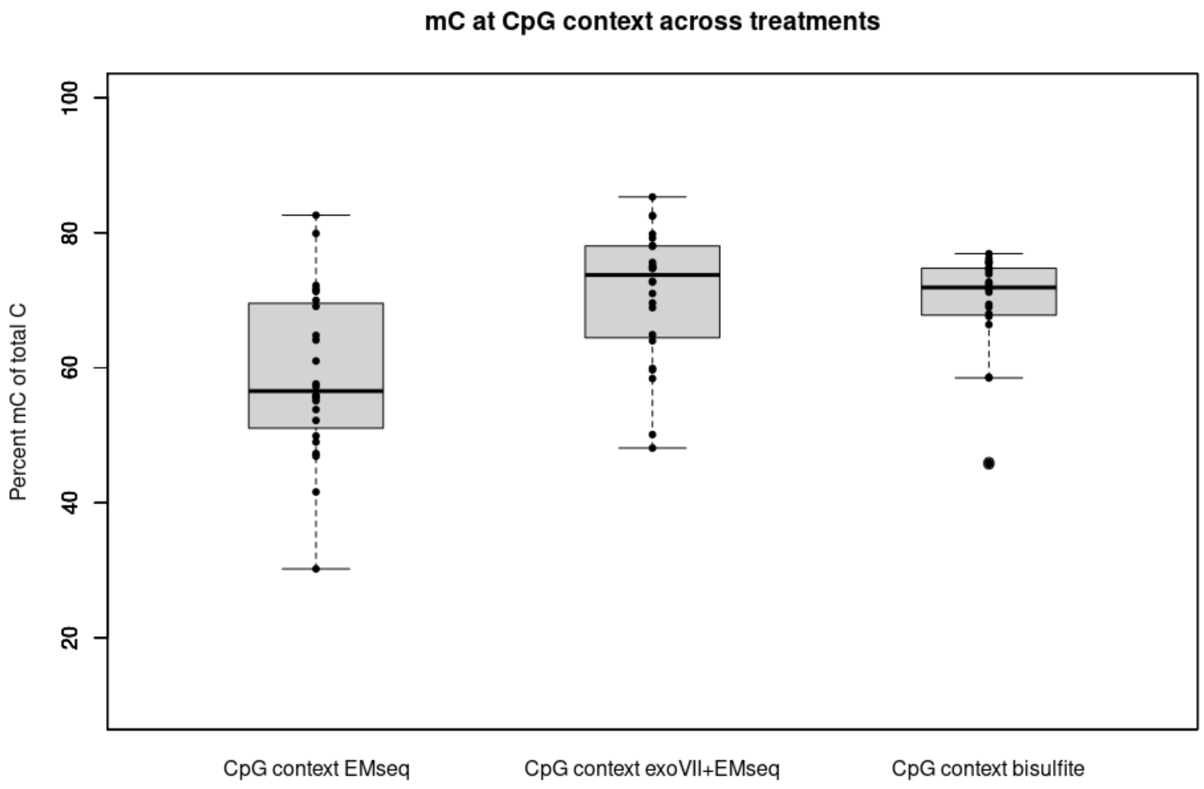


Figure S7. The percentage of methylated Cs of all Cs in CpG context of all samples used in this study comparing the EMseq method with and without exonuclease VII treatment as well as bisulfite treatment in combination with the singles stranded library method. This comparison is restricted to CpG contexts.

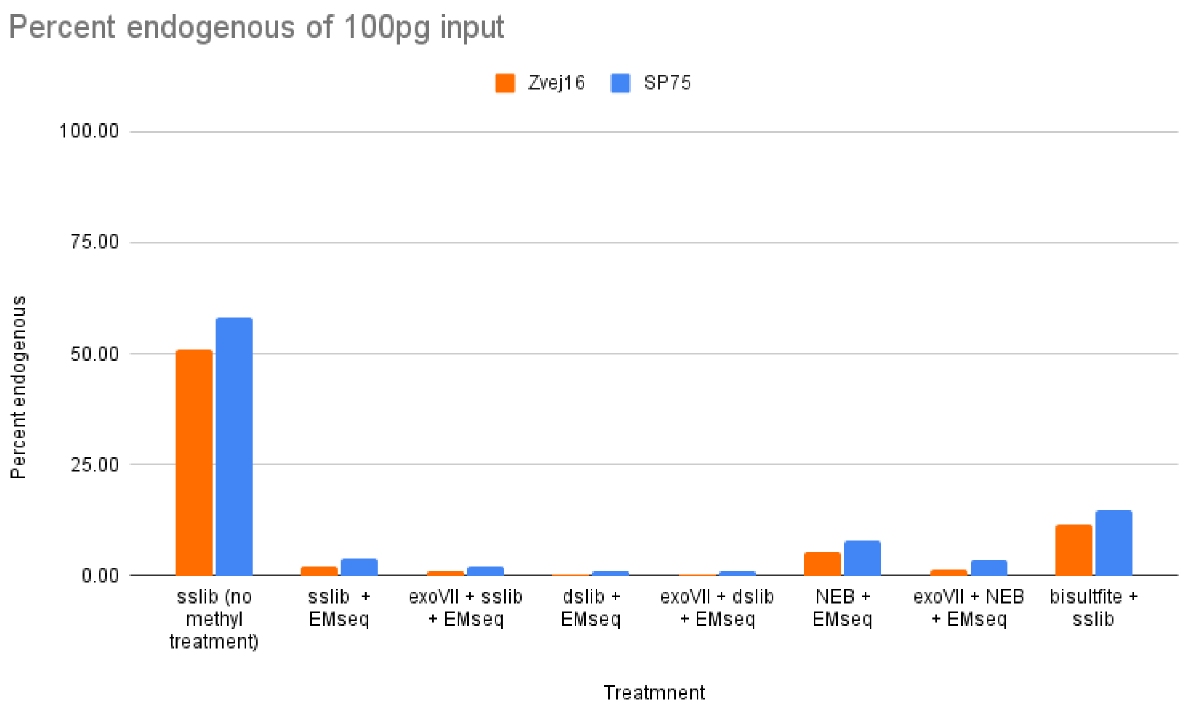


Figure S8. Percent endogenous of Zvej16 and SP75 of various treatments with 100pg input.

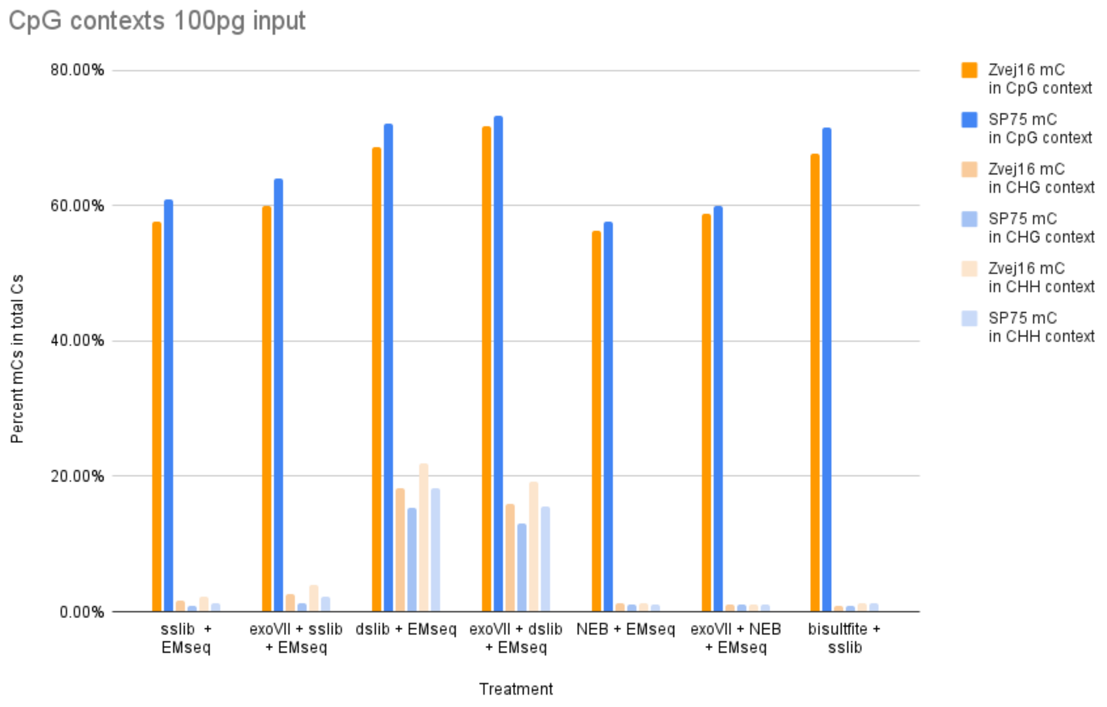


Figure S9. Percentage of mCs in different C-contexts of Zvej16 and SP75 of various treatments with 100pg input.

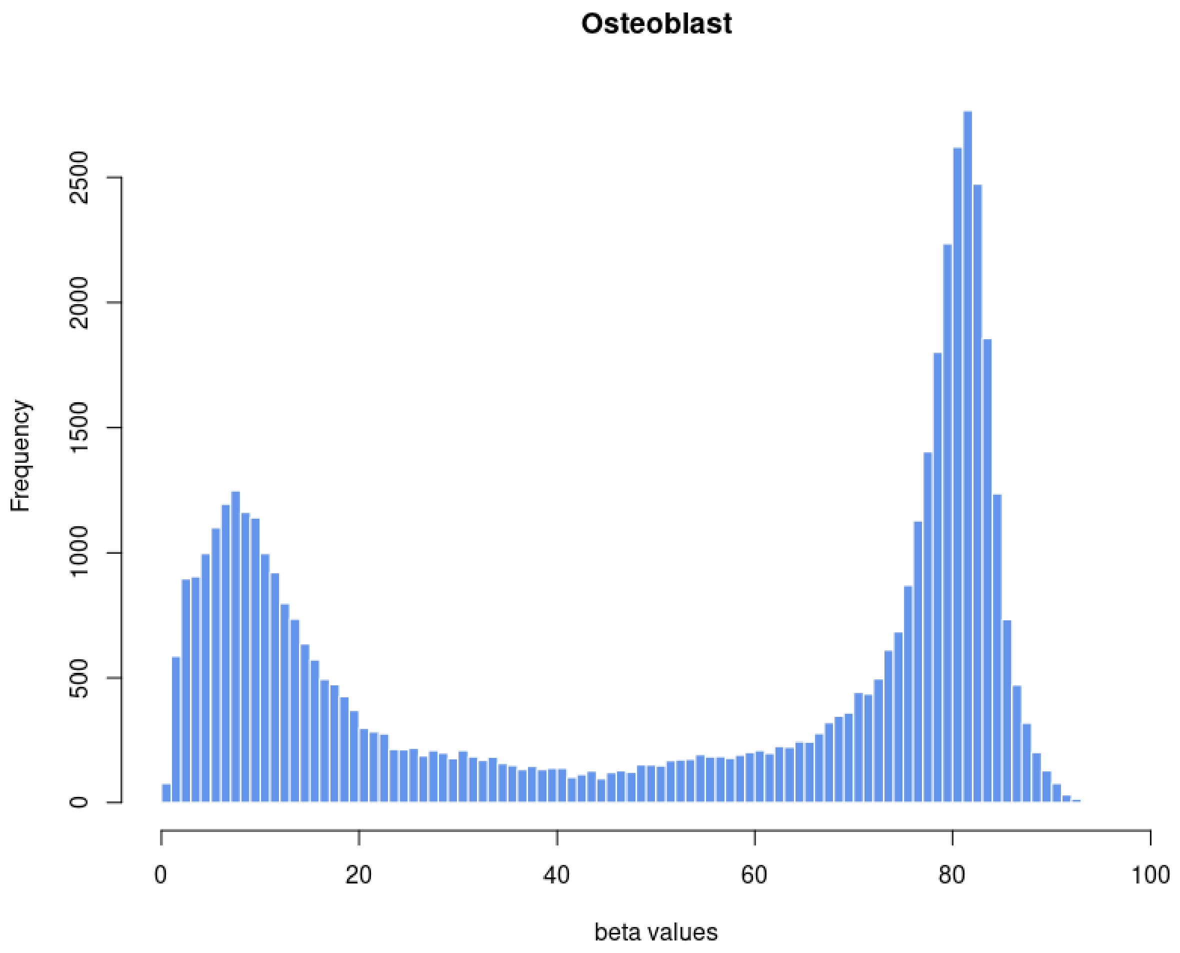


Figure S10. Histogram of segmented beta values of chromosome 1 of the high coverage osteoblast data.

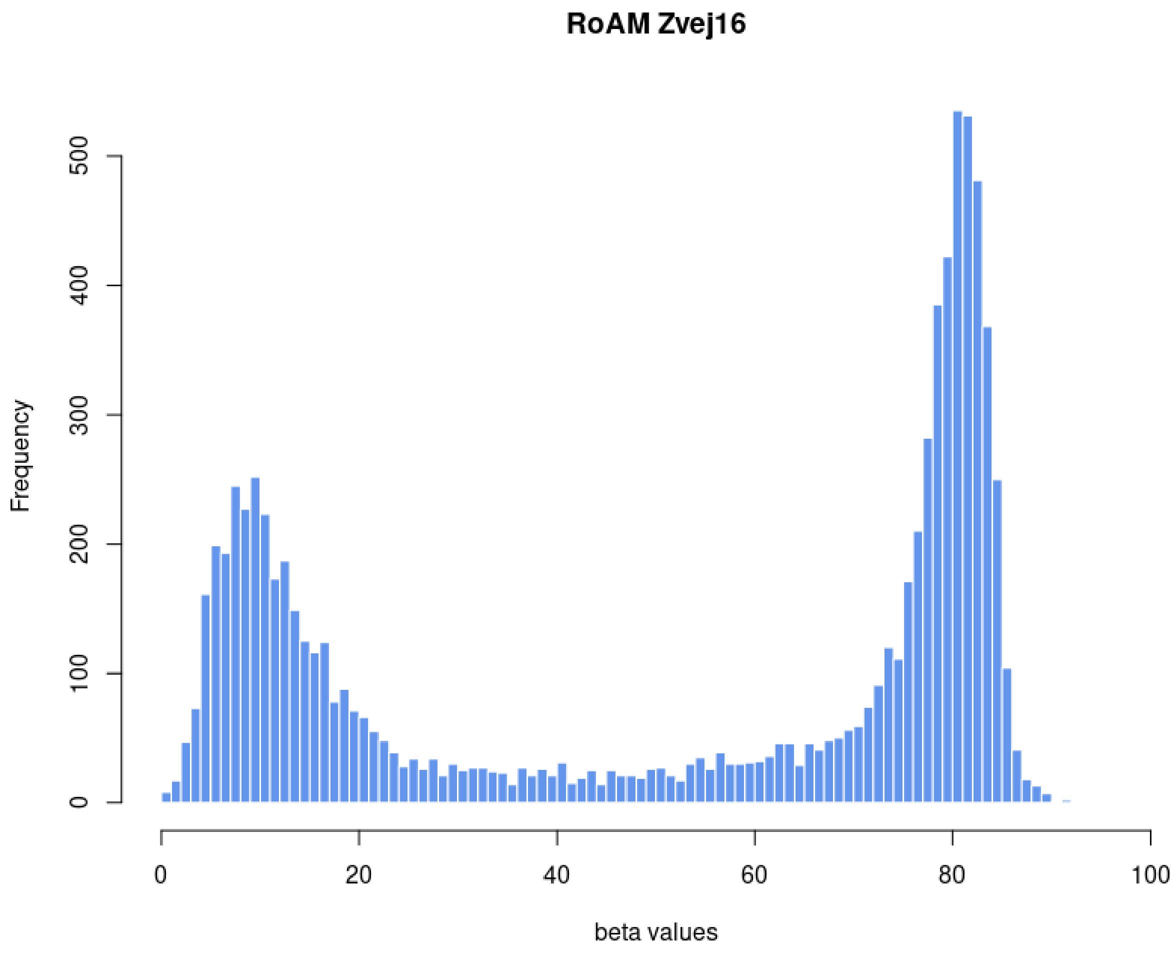


Figure S11. Histogram of segmented beta values of chromosome 1 of the 28x-fold data of Zvej16 after inferencing beta using RoAM.

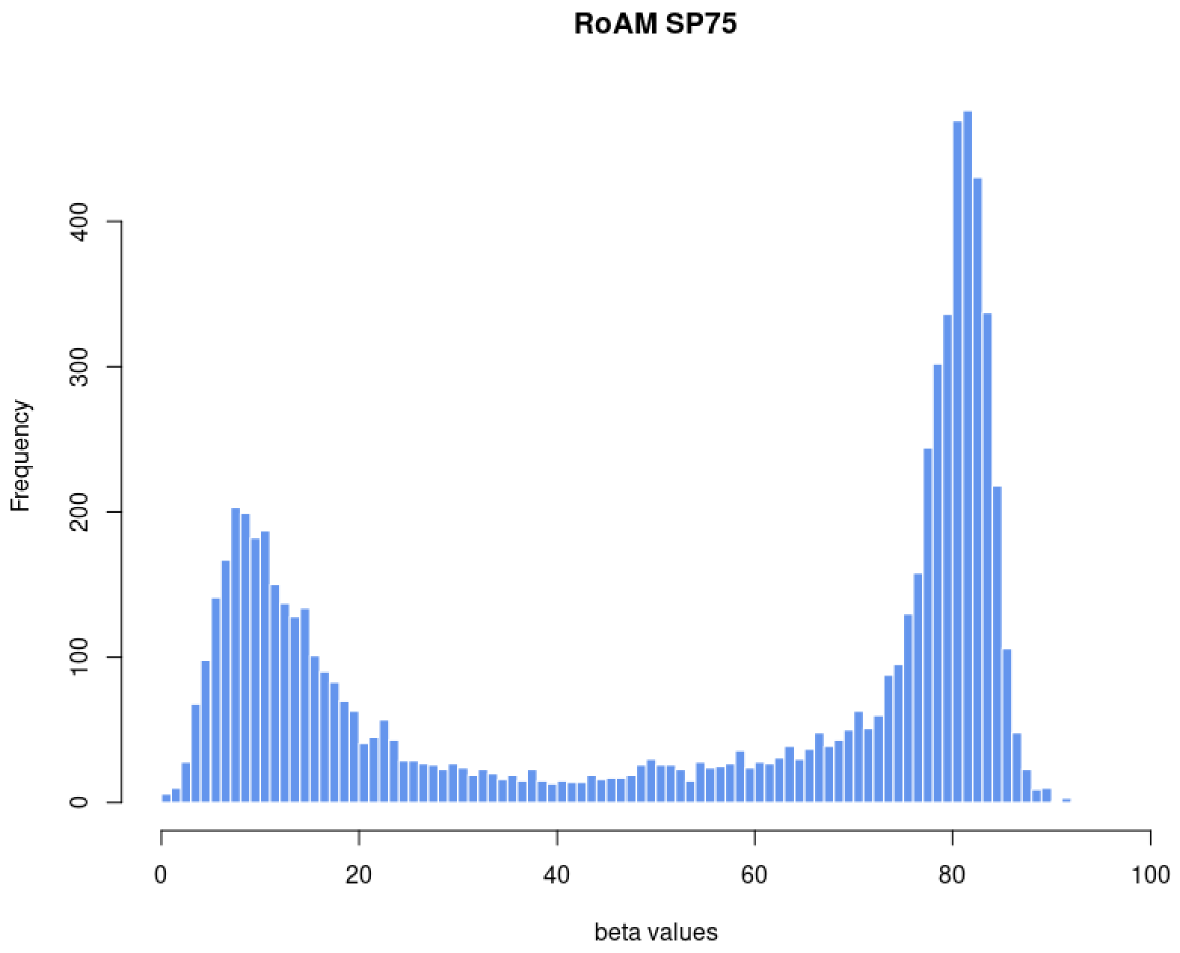


Figure S12. Histogram of segmented beta values of chromosome 1 of the 28x-fold data of SP75 after inferencing beta using RoAM.

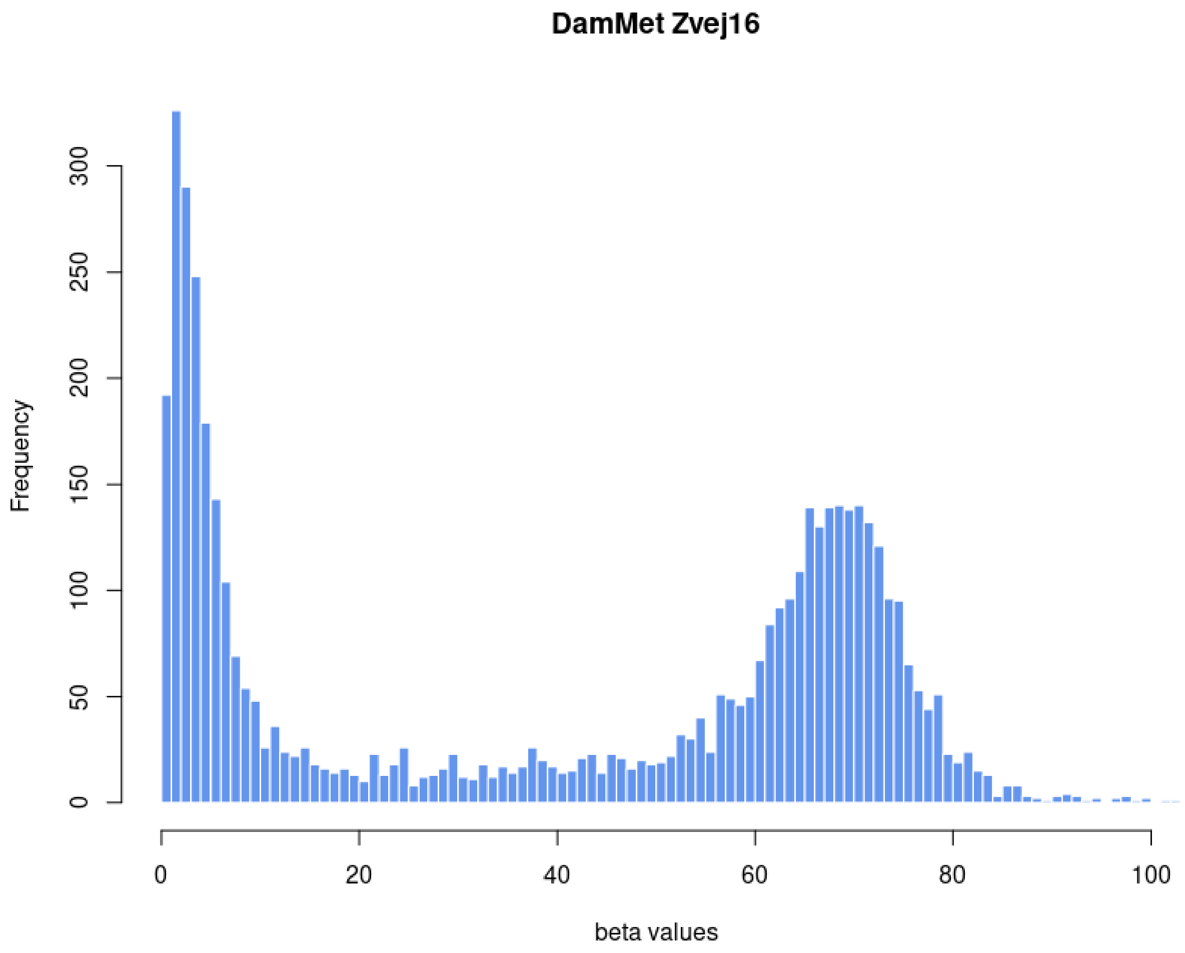


Figure S13. Histogram of segmented beta values of chromosome 1 of the 28x-fold data of Zvej16 after inferencing beta using DamMet.

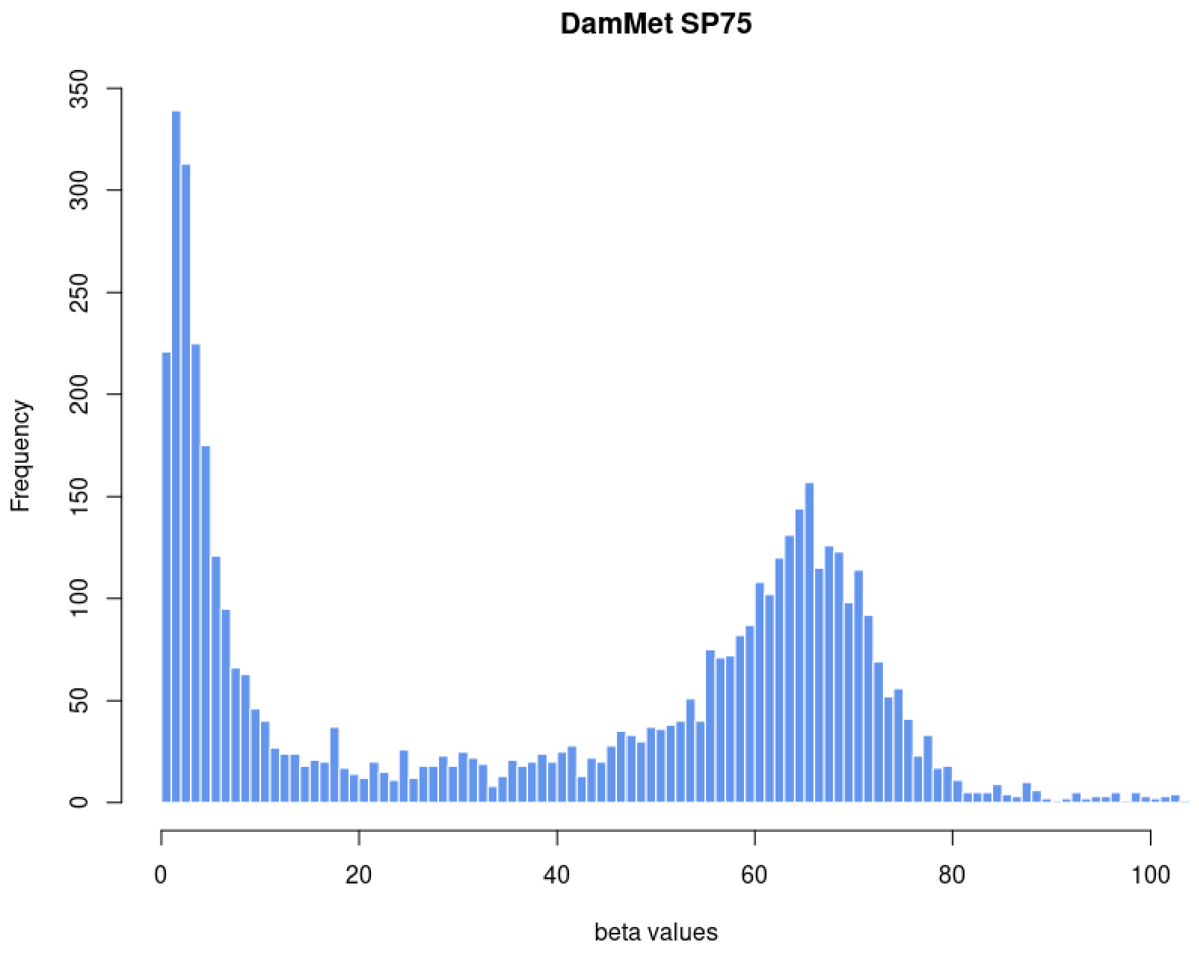


Figure S14. Histogram of segmented beta values of chromosome 1 of the 28x-fold data of SP75 after inferencing beta using DamMet.

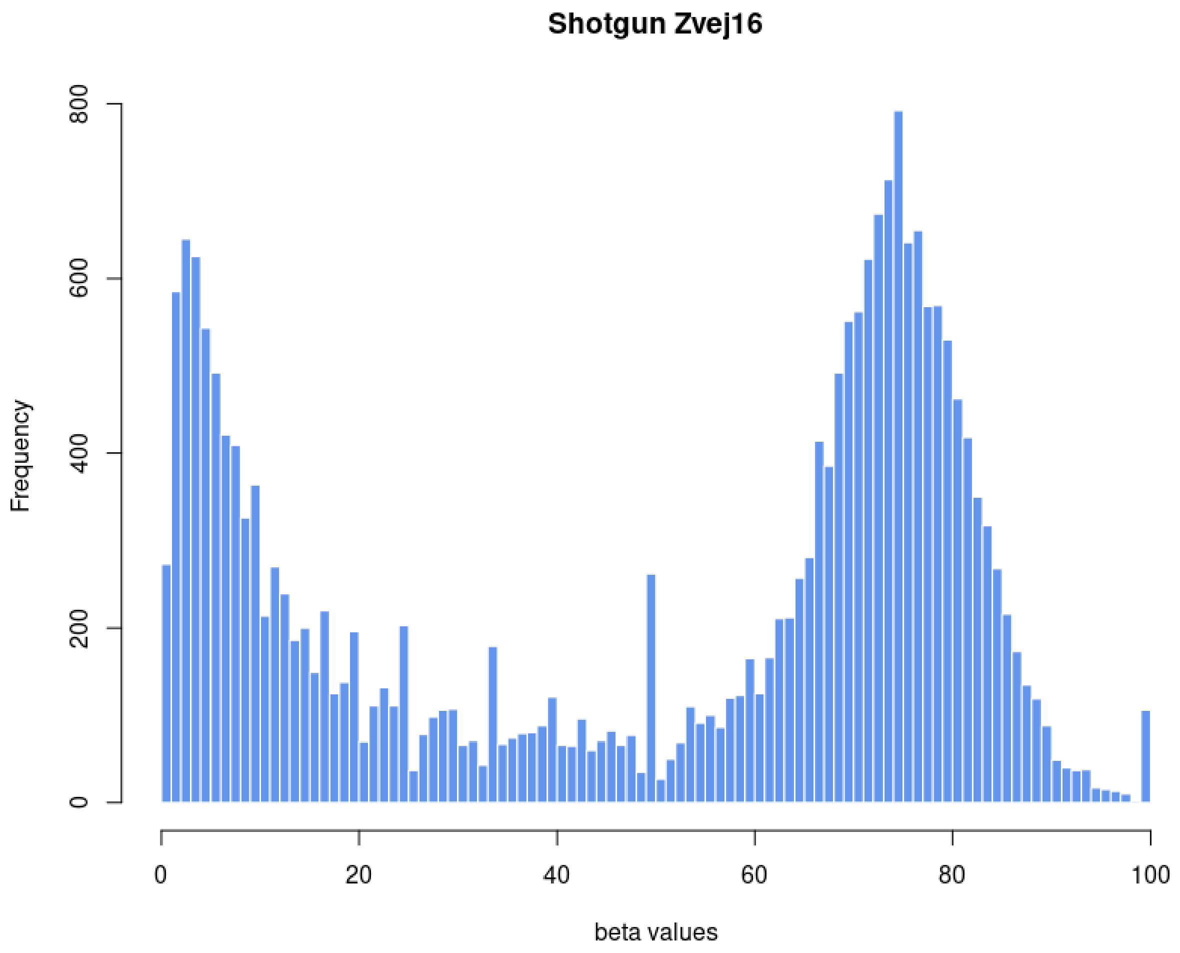


Figure S15. Histogram of segmented beta values of chromosome 1 of Zvej16 after bisulfite treatment. Beta values were calculated from shotgun data from 0.27-fold coverage.

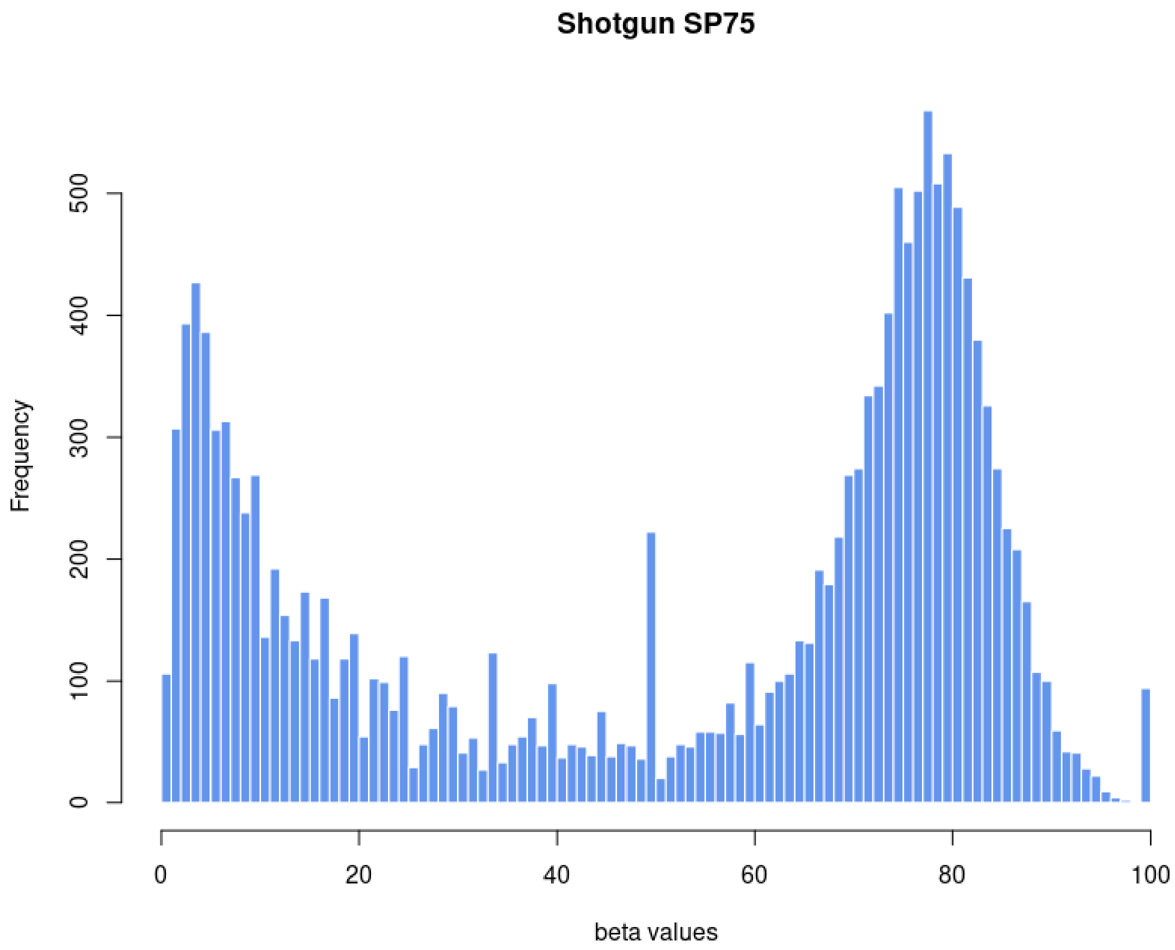


Figure S16. Histogram of segmented beta values of chromosome 1 of SP75 after bisulfite treatment. Beta values were calculated from shotgun data from 0.29-fold coverage.

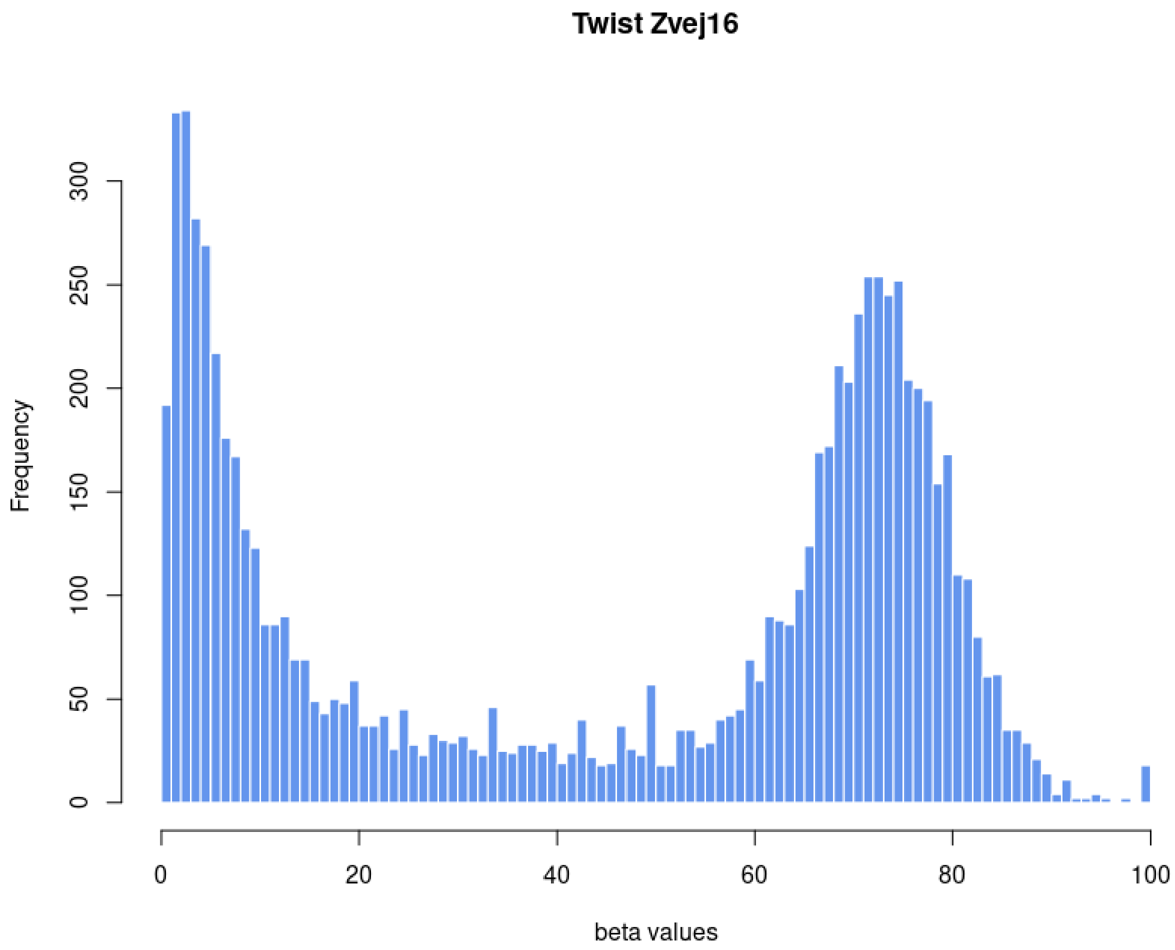


Figure S17. Histogram of segmented beta values of chromosome 1 of Zvej16 after bisulfite treatment and methylome capture using the Twist capture system.

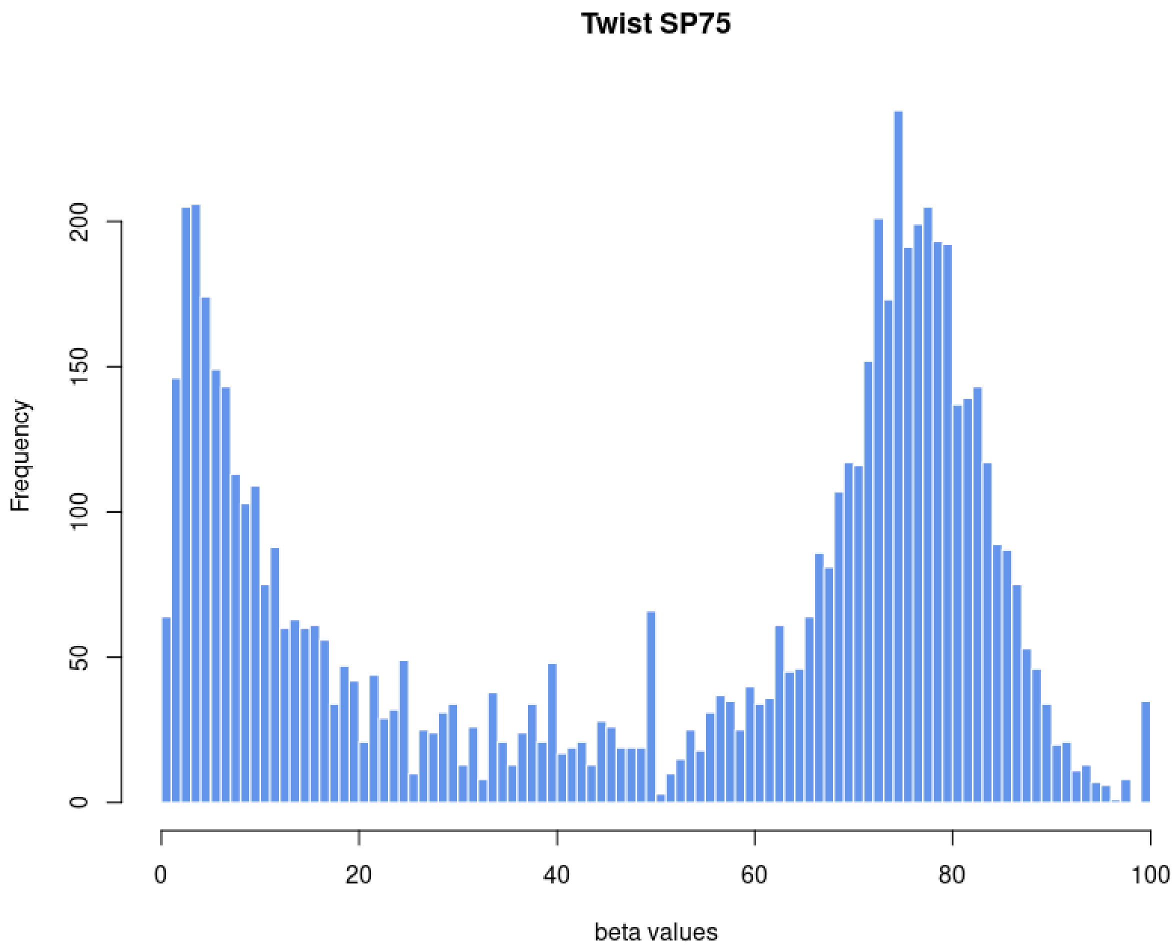


Figure S18. Histogram of segmented beta values of chromosome 1 of SP75 after bisulfite treatment and methylome capture using the Twist capture system.

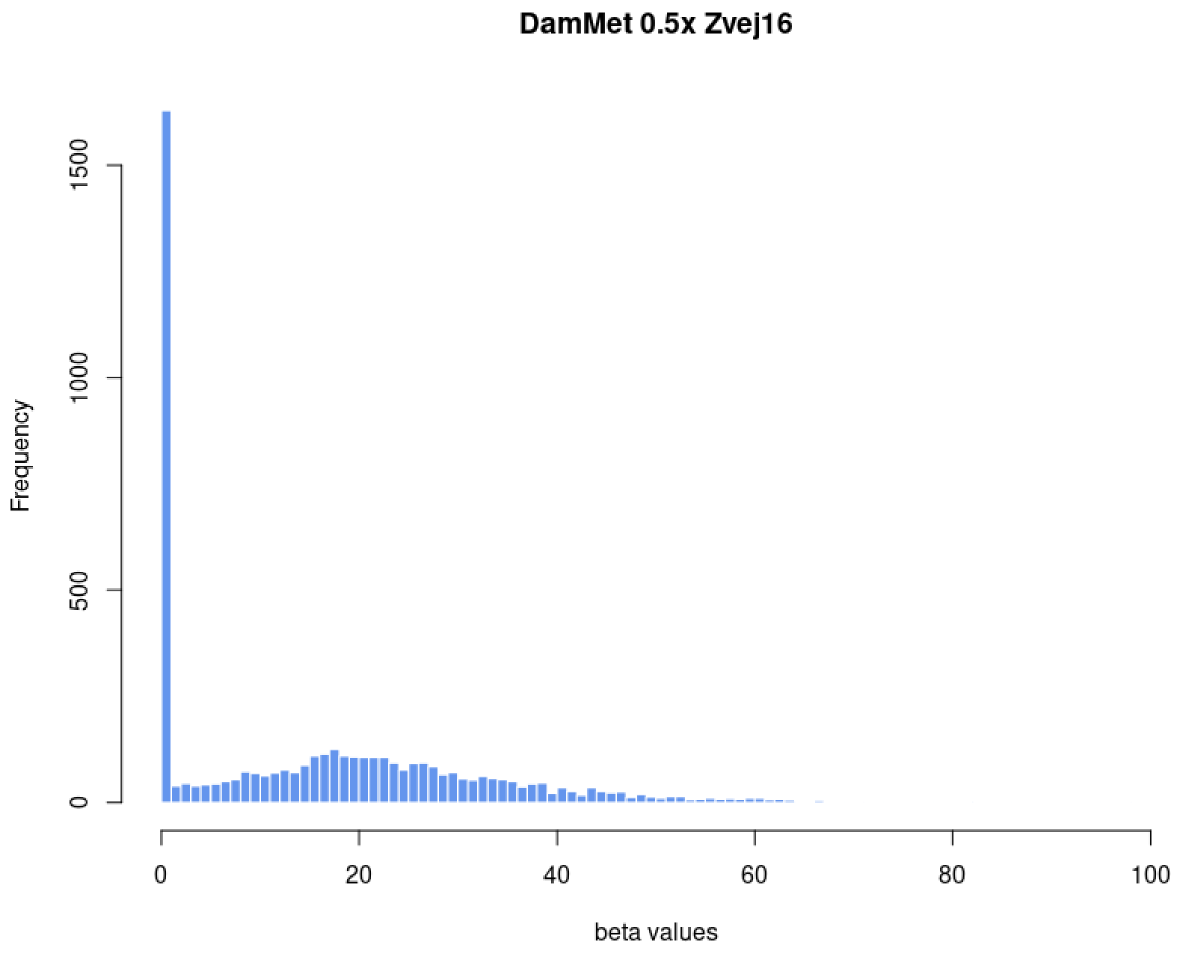


Figure S19. Histogram of segmented beta values of chromosome 1 of Zvej16 after downsampling the 28-fold non-methylation treated shotgun data to 0.5-fold coverage. Beta values were inferred using DamMet.

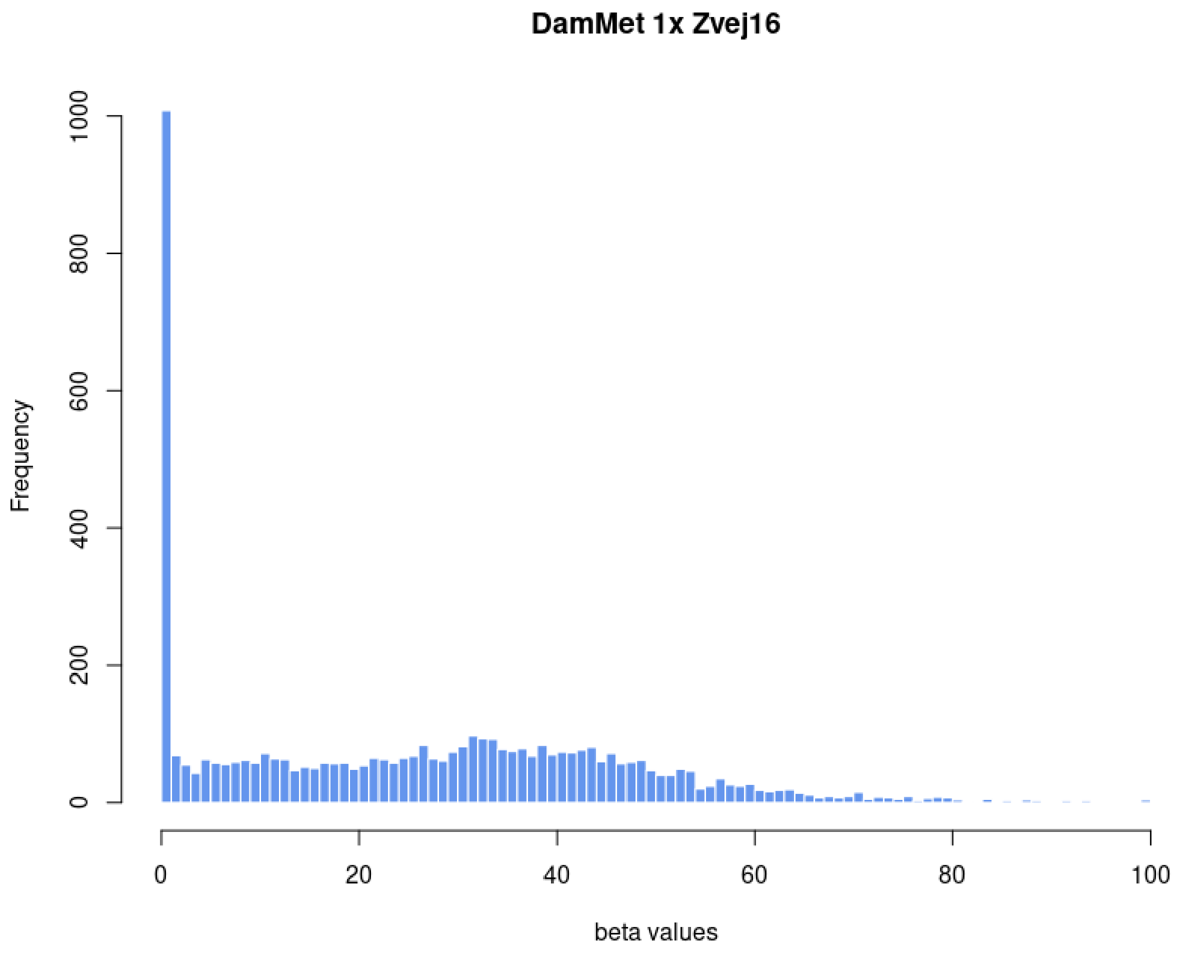


Figure S20. Histogram of segmented beta values of chromosome 1 of Zvej16 after downsampling the 28-fold non-methylation treated shotgun data to 1-fold coverage. Beta values were inferred using DamMet.

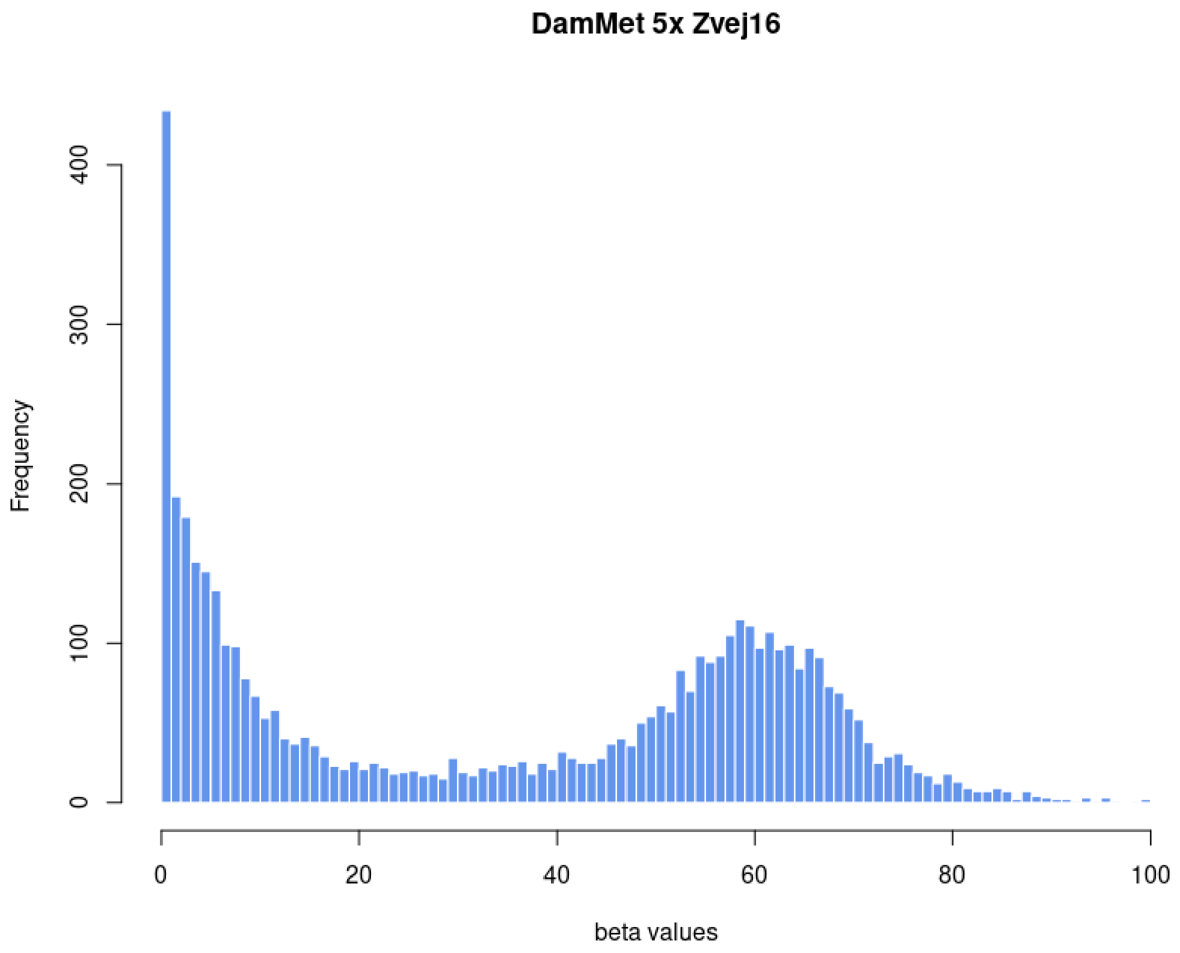


Figure S21. Histogram of segmented beta values of chromosome 1 of Zvej16 after downsampling the 28-fold non-methylation treated shotgun data to 5-fold coverage. Beta values were inferred using DamMet.

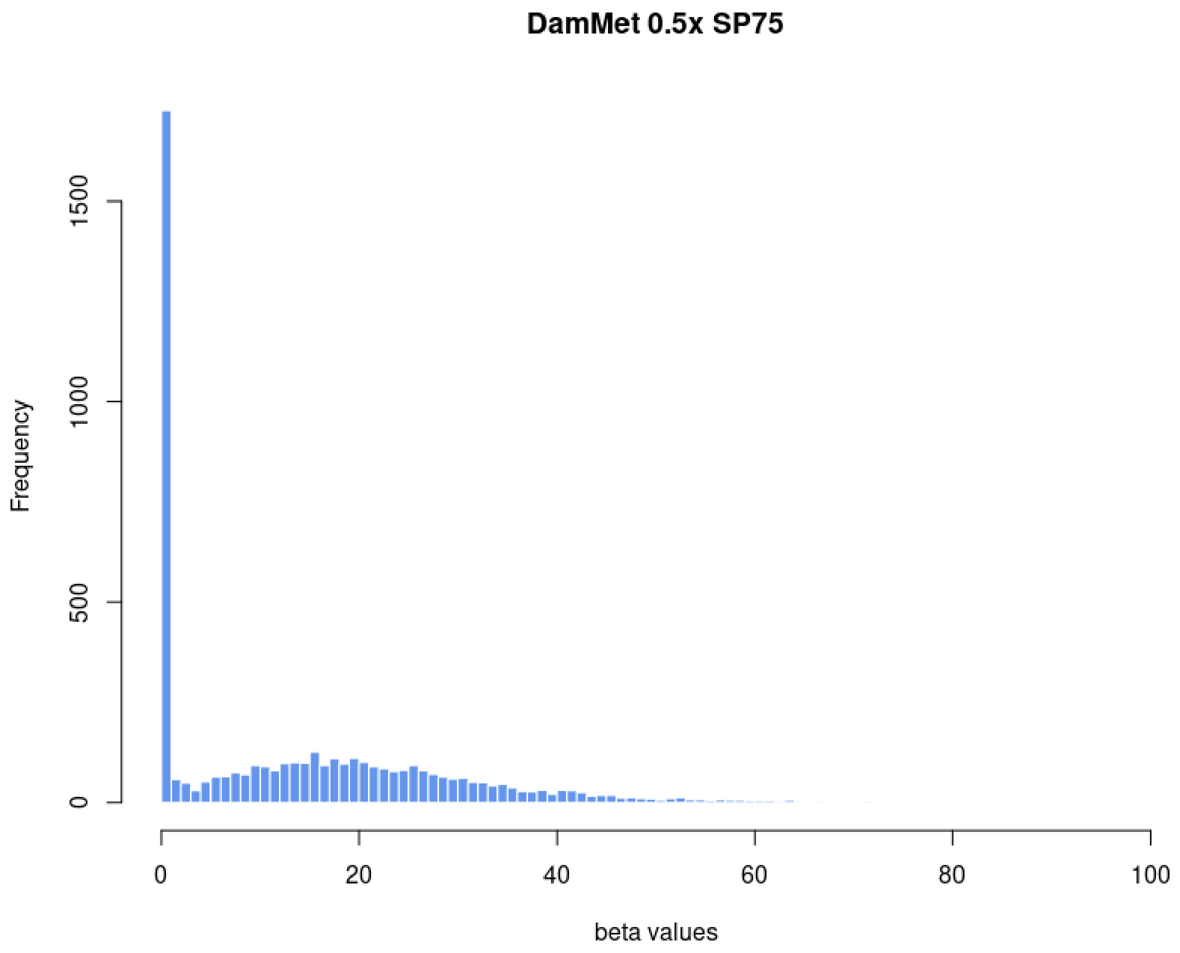


Figure S22. Histogram of segmented beta values of chromosome 1 of SP75 after downsampling the 28-fold non-methylation treated shotgun data to 0.5-fold coverage. Beta values were inferred using DamMet.

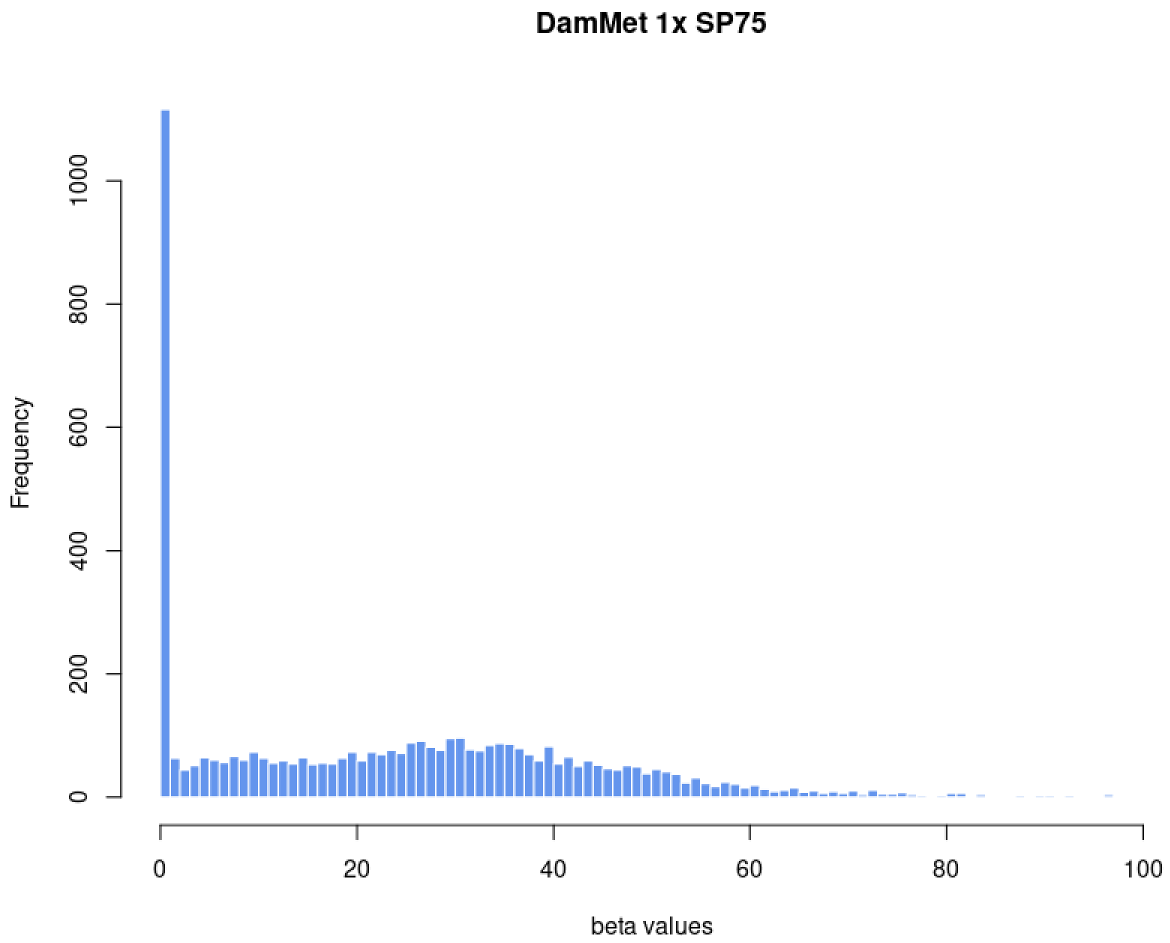


Figure S23. Histogram of segmented beta values of chromosome 1 of SP75 after downsampling the 28-fold non-methylation treated shotgun data to 1-fold coverage. Beta values were inferred using DamMet.

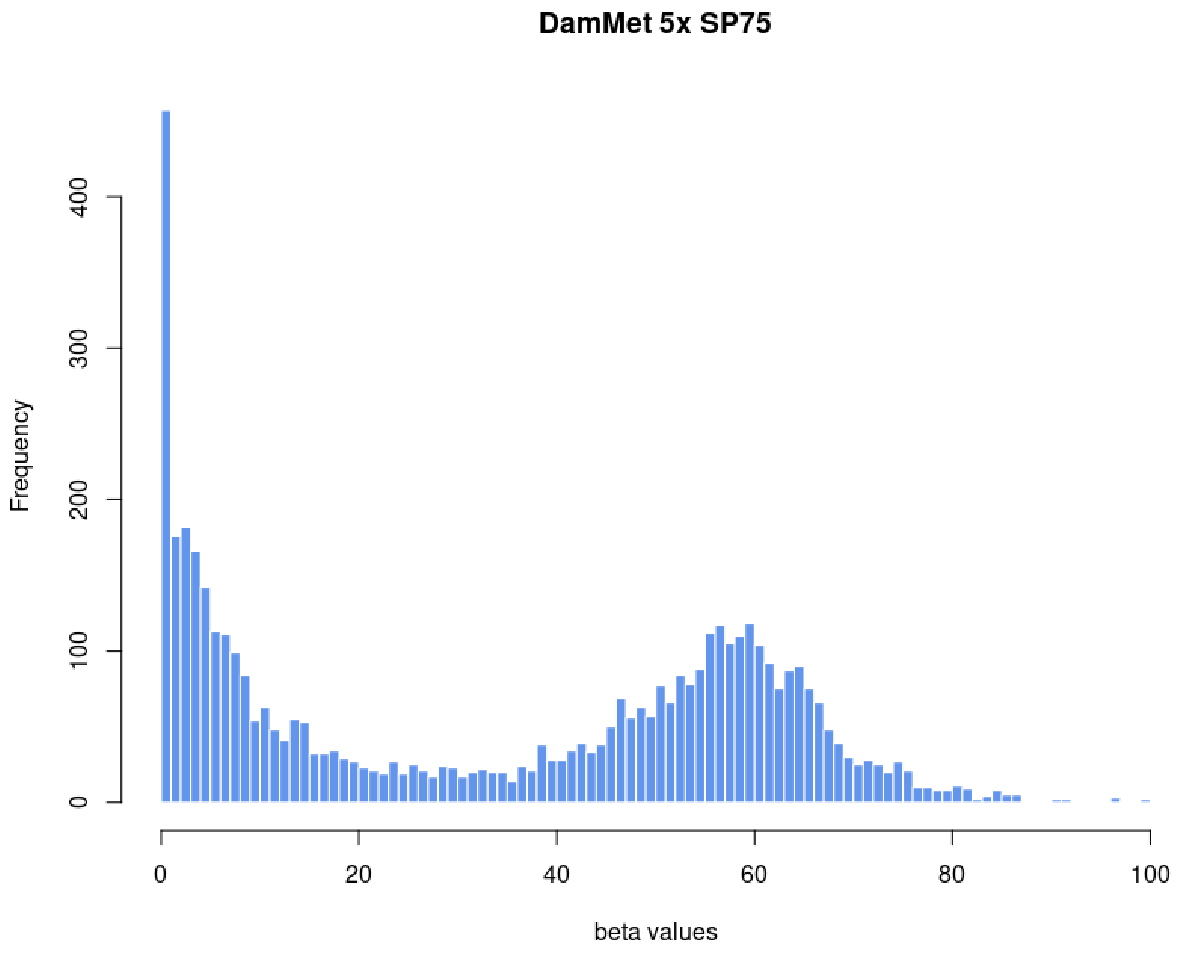


Figure S24. Histogram of segmented beta values of chromosome 1 of SP75 after downsampling the 28-fold non-methylation treated shotgun data to 5-fold coverage. Beta values were inferred using DamMet.

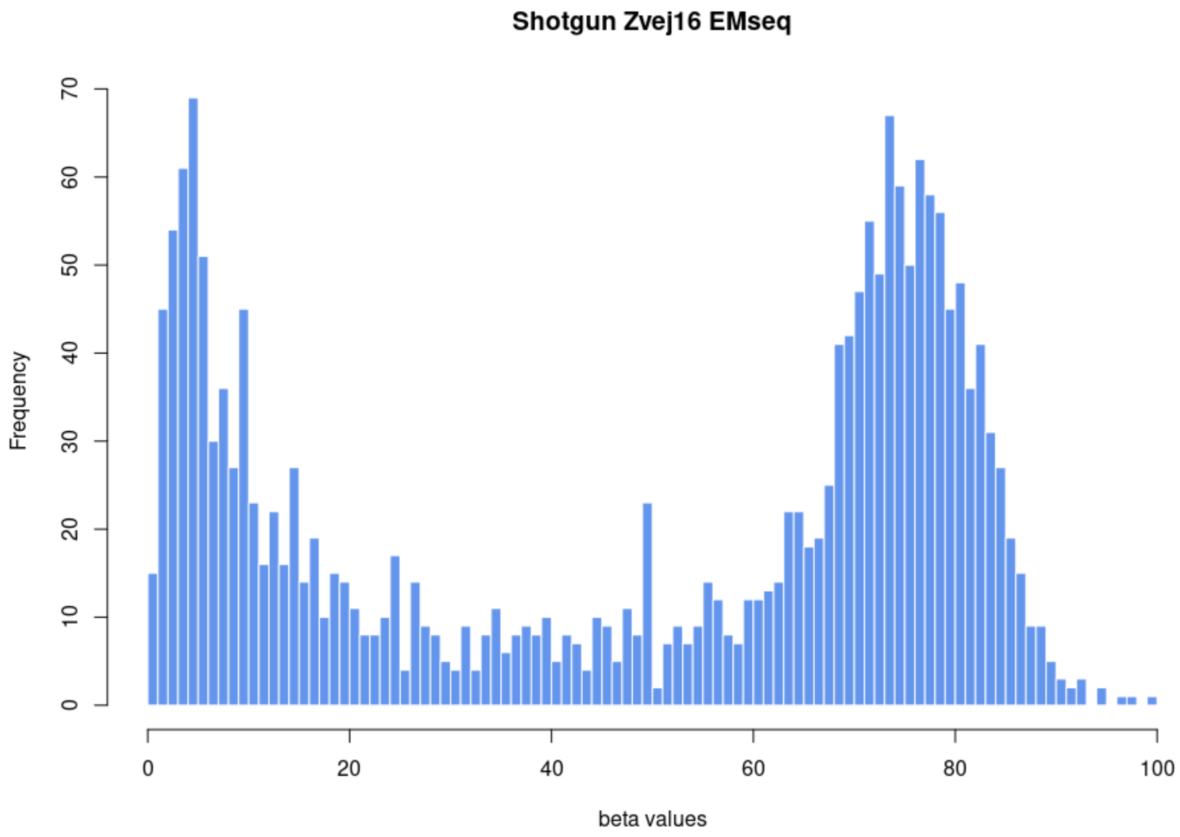


Figure S25. Histogram of segmented beta values of chromosome 1 of Zvej16 after EMseq treatment. The sample was shotgun sequenced to 0.46-fold coverage.

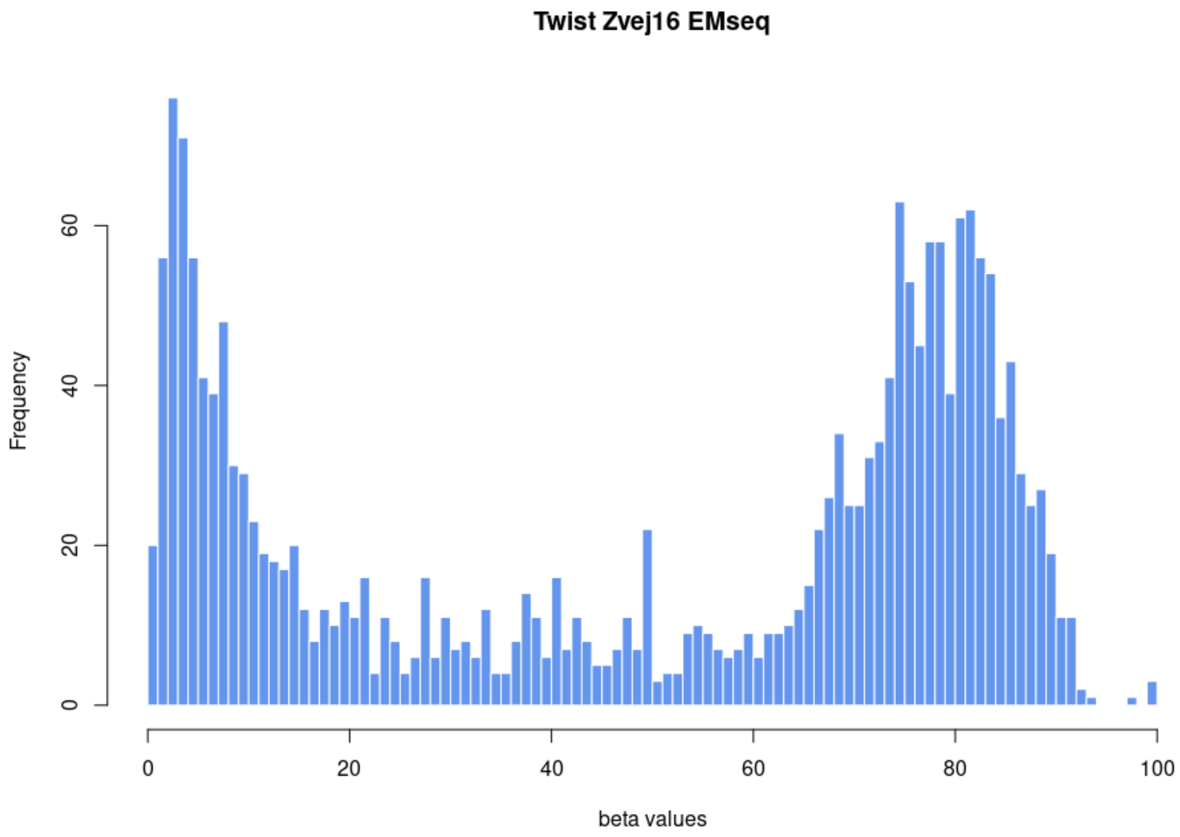


Figure S26. Histogram of segmented beta values of chromosome 1 of Zvej16 after EMseq treatment. The sample was sequenced after methylome capture using the Twist capture protocol.

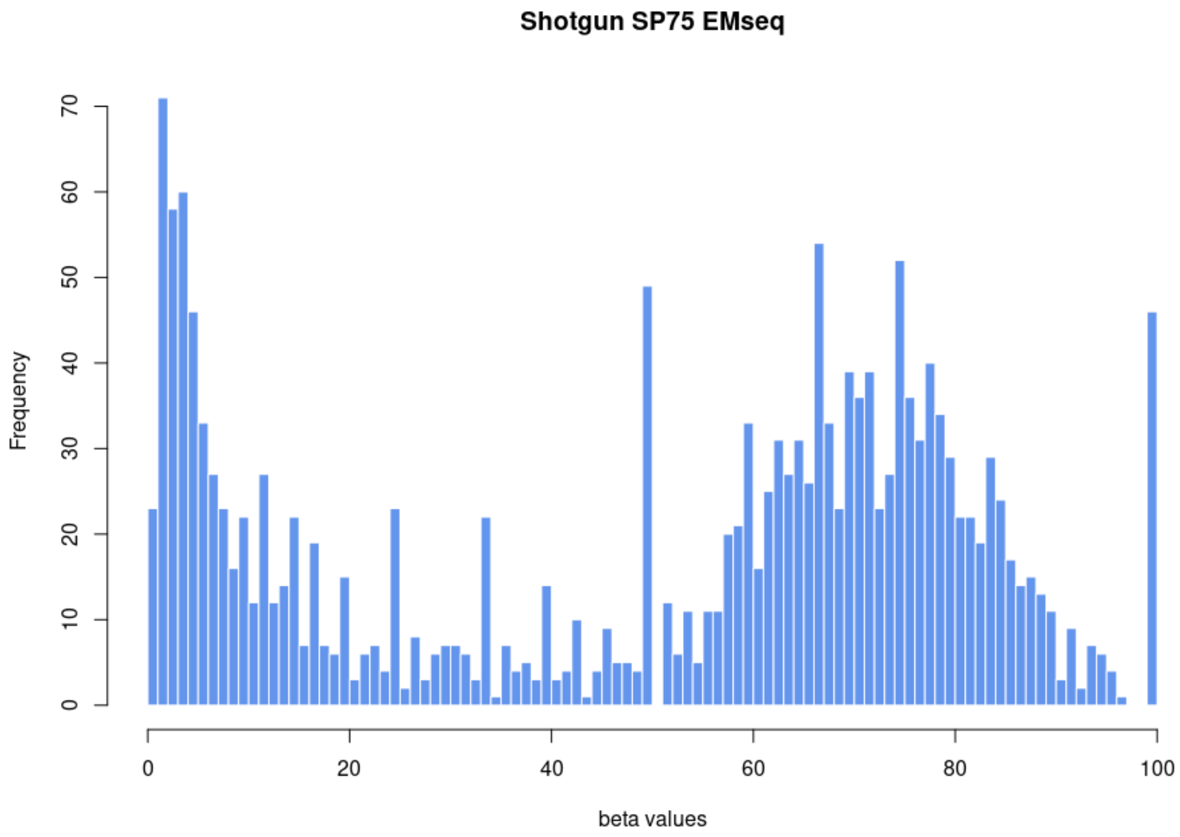


Figure S27. Histogram of segmented beta values of chromosome 1 of SP75 after EMseq treatment. The sample was shotgun sequenced to 0.61-fold coverage.

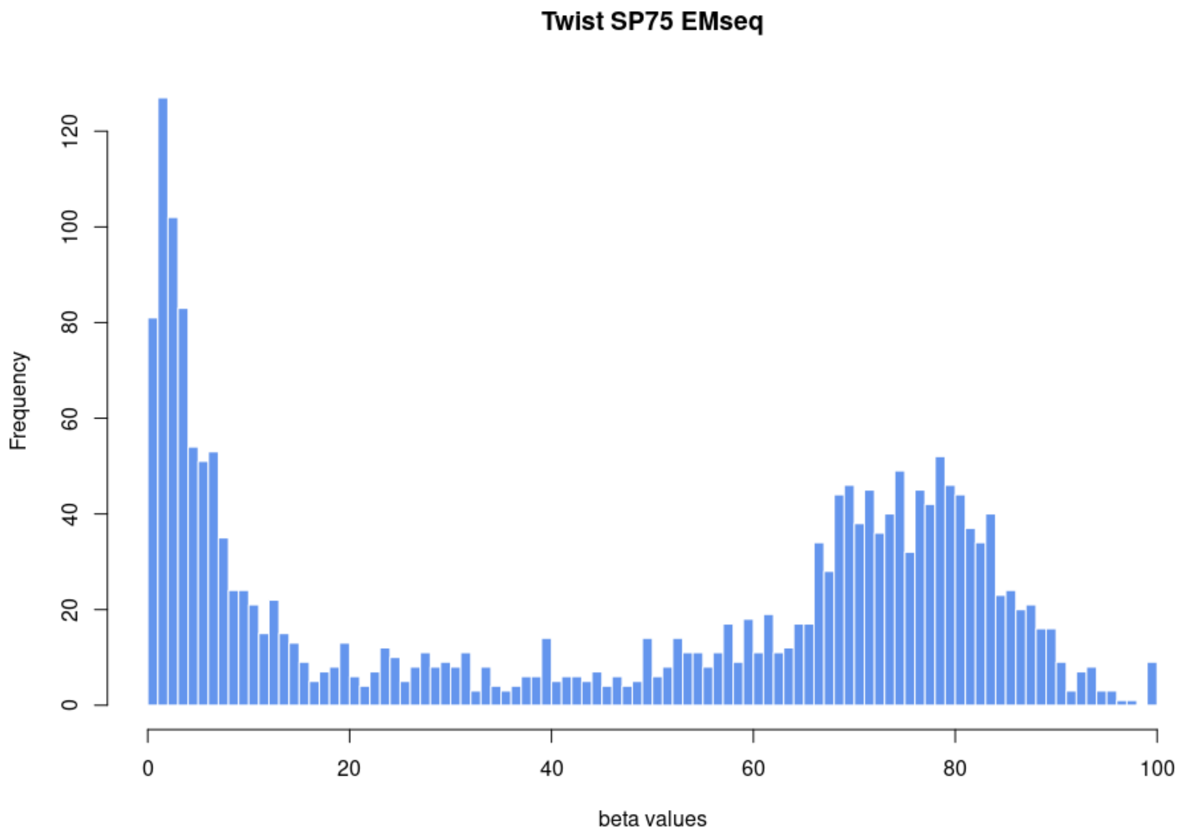


Figure S28. Histogram of segmented beta values of chromosome 1 of SP75 after EMseq treatment. The sample was sequenced after methylome capture using the Twist capture protocol.

12. Szikossy I., Kustár Á., Guba ZS., Kristóf L.A. & Pap I. Mummies from Hungary. Naturally Mummified Corpses from the Dominican Church in Vác, Hungary. In: Wieczorek A & Rosendahl, editor. Mummies of the World American Exhibitions Inc,. 160-171: Reiss-Engelhorn-Museum, Mannheim, Prestel Verl., Munich/Berlin/London/New York,;

13. Szikossy I. KLA&. PI. Mummies found in the Dominican Church of Vác, Hungary. In: Wieczorek A & Rosendhal, editor. Mummies of the World American Exhibitions Inc,. Reiss-Engelhorn-Museum, Mannheim, Prestel Verl., Munich/Berlin/London/New York; pp. 355–356.

14. Donoghue HD, Pap I, Szikossy I, Spigelman M. Detection and characterization of Mycobacterium tuberculosis DNA in 18th Century Hungarians with pulmonary and extra-pulmonary tuberculosis. In: Gill-Frerking, H and Rosendahl, W and Zink, A and Piombino-Mascasli, D, editor. Yearbook of mummy studies. Verlag Dr. Friedrich Pfeil: München, Germany; 2011. pp. 51–56.

15. Donoghue HD, Pap I, Szikossy I, Spigelman M. The Vác Mummy Project: Investigation of 265 Eighteenth-Century Mummified Remains from the TB Pandemic Era. In: Shin DH, Bianucci R, editors. The Handbook of Mummy Studies: New Frontiers in Scientific and Cultural Perspectives. Singapore: Springer Singapore; 2021. pp. 777–805.

16. Pinhasi R, Fernandes DM, Sirak K, Cheronet O. Isolating the human cochlea to generate bone powder for ancient DNA analysis. Nat Protoc. 2019;14: 1194–1205.

17. Dabney J, Knapp M, Glocke I, Gansauge M-T, Weihmann A, Nickel B, et al. Complete mitochondrial genome sequence of a Middle Pleistocene cave bear reconstructed from ultrashort DNA fragments. Proc Natl Acad Sci U S A. 2013;110: 15758–15763.

18. Meyer M, Kircher M. Illumina sequencing library preparation for highly multiplexed target capture and sequencing. Cold Spring Harb Protoc. 2010;2010: db.prot5448.

25. R. Core Team. An Introduction to R. Samurai Media Limited; 2015.

26. Li H, Durbin R. Fast and accurate short read alignment with Burrows-Wheeler transform. Bioinformatics. 2009;25: 1754–1760.

27. Loyfer N, Magenheim J, Peretz A, Cann G, Bredno J, Klochendler A, et al. A DNA methylation atlas of normal human cell types. Nature. 2023;613: 355–364.

28. Gokhman D, Nissim-Rafinia M, Agranat-Tamir L, Housman G, García-Pérez R, Lizano E, et al. Differential DNA methylation of vocal and facial anatomy genes in modern humans. Nat Commun. 2020;11: 1189.

29. Hanghøj K, Renaud G, Albrechtsen A, Orlando L. DamMet: ancient methylome mapping

accounting for errors, true variants, and post-mortem DNA damage. Gigascience. 2019;8. doi:10.1093/gigascience/giz025

30. Mathieson I, Alpaslan-Roodenberg S, Posth C, Szécsényi-Nagy A, Rohland N, Mallick S, et al. The genomic history of southeastern Europe. Nature. 2018;555: 197–203.

31. Zeng TC, Vyazov LA, Kim A, Flegontov P, Sirak K, Maier R, et al. Postglacial genomes from foragers across Northern Eurasia reveal prehistoric mobility associated with the spread of the Uralic and Yeniseian languages. bioRxiv. 2023. doi:10.1101/2023.10.01.560332

32. Quinlan AR, Hall IM. BEDTools: a flexible suite of utilities for comparing genomic features. Bioinformatics. 2010;26: 841–842.

33. Akalin A, Kormaksson M, Li S, Garrett-Bakelman FE, Figueroa ME, Melnick A, et al. methylKit: a comprehensive R package for the analysis of genome-wide DNA methylation profiles. Genome Biol. 2012;13: R87.
